# An improved auxin-inducible degron system for fission yeast

**DOI:** 10.1101/2021.07.19.452993

**Authors:** Xiao-Ran Zhang, Lei Zhao, Fang Suo, Yadong Gao, Qingcui Wu, Xiangbing Qi, Li-Lin Du

**Author notes:** Corresponding author: National Institute of Biological Sciences, 7 Science Park Road, Zhongguancun Life Science Park, Beijing 102206, China.

## Abstract

Conditional degron technologies, which allow a protein of interest to be degraded in an inducible manner, are important tools for biological research, and are especially useful for creating conditional loss-of-function mutants of essential genes. The auxin-inducible degron (AID) technology, which utilizes plant auxin signaling components to control protein degradation in non-plant species, is a widely used small-molecular-controlled degradation method in yeasts and animals. However, the currently available AID systems still have room for further optimization. Here, we have improved the AID system for the fission yeast *Schizosaccharomyces pombe* by optimizing all three components: the AID degron, the small-molecule inducer, and the inducer-responsive F-box protein. We chose a 36-amino-acid sequence of the *Arabidopsis* IAA17 protein as the degron and employed three tandem copies of it to enhance efficiency. To minimize undesirable side effects of the inducer, we adopted a bulky analog of auxin, 5-adamantyl-IAA, and paired it with the F-box protein OsTIR1 that harbors a mutation (F74A) at the auxin-binding pocket. 5-adamantyl-IAA, when utilized with OsTIR1-F74A, is effective at concentrations thousands of times lower than auxin used in combination with wild-type OsTIR1. We tested our improved AID system on 10 essential genes and achieved inducible lethality for all of them, including ones that could not be effectively inactivated using a previously published AID system. Our improved AID system should facilitate the construction of conditional loss-of-function mutants in fission yeast.

## INTRODUCTION

Studies of gene functions, especially the functions of essential genes, are greatly assisted by the conditional inactivation of genes. Diverse strategies have been employed to achieve conditional disruption of gene functions. For model yeast species, these strategies include creating temperature-sensitive mutants (Dohmen et al. 1994; Kanemaki et al. 2003; Ben-Aroya et al. 2008), placing a target gene under the control of a repressible promoter (Mnaimneh et al. 2004), and chemically induced alteration of the localization of the gene product (Haruki et al. 2008). One particularly powerful strategy that has gained wide-spread use in recent years is the auxin-inducible degron (AID) technology, which adopts a protein degradation mechanism in plants for conditional depletion of proteins in non-plant organisms (Nishimura et al. 2009; Kanke et al. 2011; Nishimura and Kanemaki 2014; Zhang et al. 2015; Natsume et al. 2016; Natsume and Kanemaki 2017; Texier et al. 2018).

In plants, auxin, a plant hormone, promotes the degradation of auxin/indole-3-acetic acid proteins (Aux/IAAs) by acting as a molecular glue to mediate the interactions between Aux/IAAs and transport inhibitor response 1/auxin signaling F-box proteins (TIR1/AFBs), which are F-box proteins in SCF (Skp1, Cullin, and F-box protein) E3 ubiquitin ligase complexes (Tan et al. 2007). Aux/IAAs and TIR1/AFBs only exist in land plants (Mutte et al. 2018), but when an Aux/IAA protein and a TIR1/AFB protein are introduced together into yeast or animal cells, they are sufficient to recapitulate auxin-induced degradation by acting with endogenous SCF components (Nishimura et al. 2009). This is the mechanistic basis of the AID technology.

An AID system is composed of three components: a degron fused to the target protein, an F-box protein, and a small molecule inducer. The IAA17 protein from *Arabidopsis thaliana*, which has a short half-life in seedlings (Dreher et al. 2006), has often been used as the degron (Nishimura et al. 2009; Kanke et al. 2011). The most widely used F-box proteins in AID systems are the TIR1 proteins from *Arabidopsis thaliana* and rice (*Oryza sativa*), which are referred to as AtTIR1 and OsTIR1, respectively. OsTIR1 works better than AtTIR1 at temperatures higher than 24°C and is a preferred choice for yeasts and mammalian cells, whose optimal growth temperatures are higher than 24°C (Nishimura et al. 2009).

The widely used small molecule inducer in AID systems, the auxin indole-3-acetic acid (IAA) or its synthetic analog naphthalene acetic acid (NAA), is generally assumed to be inert in non-plant species. However, over 50 years ago, physiological effects of IAA and NAA on certain yeast species have been described (Yanagishima and Masuda 1965; Kamisaka et al. 1967). Many fungal species are now known to synthesize auxin and/or exhibit physiological responses to auxin (Fu et al. 2015; Chanclud and Morel 2016). In particular, the model organism *Saccharomyces cerevisiae* is able to synthesize and secrete auxin (Rao et al. 2010). Furthermore, in *S. cerevisiae*, IAA promotes the morphological transition to a filamentous form, retards growth, and inhibits the activity of TORC1 (Prusty et al. 2004; Liu et al. 2016; Snyder et al. 2019; Nicastro et al. 2021), and NAA exhibits a stronger TORC1 inhibition effect than IAA (Prusty et al. 2004; Liu et al. 2016; Snyder et al. 2019; Nicastro et al. 2021). In the other model yeast, the fission yeast *Schizosaccharomyces pombe*, growth inhibition was observed in the presence of high concentrations of NAA (Kanke et al. 2011). Thus, using IAA or NAA as an inducer in the AID systems, especially at a high concentration, may cause unintended consequences in yeasts.

The AID technology was first adopted for use in *S. pombe* by Kanke et al. (Kanke et al. 2011). In the AID system of Kanke et al. (named the *i*-AID system by Kanke et al. and hereafter referred to as the Kanke system), plant F-box proteins were fused to *S. pombe* Skp1 to increase the efficiency of target protein depletion. However, Skp1-fused AtTIR1 is toxic to *S. pombe* when expressed from a strong promoter (Kanke et al. 2011). Thus, in the Kanke system, a moderate-strength promoter, *Padh15*, was used to express the Skp1-fused F-box proteins. The Kanke system was tested on 15 essential genes. Out of these genes, severe growth inhibition was observed for only 3 genes and no effect was detected for 6 genes (Kanke et al. 2011). Combining the Kanke system with a transcription shut-off approach resulted in conditional lethality for all 15 essential genes (Kanke et al. 2011). It is desirable to further improve the efficiency of the AID system for *S. pombe* so that the AID system alone is sufficient to generate conditional null phenotypes for a high percentage of genes.

Here, we report an improved AID system for *S. pombe*. We show that our system is more efficient than the Kanke system and can generate conditional lethality for all essential genes tested, including ones that are refractory to the Kanke system.

## MATERIALS AND METHODS

### Plasmid construction

Plasmids used in this study are listed in Table 1. The C-terminal degron tagging plasmid pDB4581 was constructed by cloning a synthetic DNA fragment into a kanMX-marked pFA6a-derived C-terminal tagging plasmid backbone. The synthetic DNA fragment, which contains the sequence encoding the XTEN16 linker (SGSETPGTSESATPES) and three tandem copies of a 36-amino-acid region of AtIAA17 (amino acids 71-106 of AtIAA17, termed sAID for short AID), was synthesized by Wuxi Qinglan Biotech. When designing the DNA sequence, we preferentially chose codons favored by *S. pombe*. To reduce the chance of recombination between the three copies of sAID-coding sequences, codons were chosen so that between-copy nucleotide identities are lower than 80%.

**Table 1.** The plasmids created and used in this study.

| Plasmid<br>number | Addgene ID | Description | Purpose |
| --- | --- | --- | --- |
| pDB4581 | 171124 | *pFA6a-3×sAID-kanMX* | 3×sAID C-terminal tagging |
| pDB4697 | 171125 | vector_ura4-integrating | No OsTIR1 control |
| pDB5050 | 171126 | *Padh1-OsTIR1_ura4-*<br>integrating | Expressing wild-type OsTIR1 (paired<br>with NAA) |
| pDB4695 | 171127 | *Padh1-OsTIR1-F74A_ura4-*<br>integrating | Expressing OsTIR1-F74A (paired with<br>5-Adamantyl-IAA) |

Plasmids for integrating the OsTIR1 expression cassettes are based on the vector plasmid pDB4697. pDB4697 was constructed by cloning a 2567-bp *ura4*-gene-containing sequence from the *S. pombe* genome (chromosome III: 114,830-117,653) into the pEASY-blunt vector (TransGen Biotech) and placing it between two NotI sites. This sequence contains the 1.8-kb region deleted in *ura4-D18* and sequences flanking the 1.8-kb deleted region (about 400 bp on each side). *Padh1-OsTIR1* integrating plasmid pDB5050 was constructed by cloning the *S. pombe Padh1* promoter, OsTIR1 coding sequence codon-optimized for *S. pombe* (synthesized by Wuxi Qinglan Biotech), and the terminator of *S. cerevisiae ADH1* gene into an AvrII site about 70 bp downstream of the coding sequence of *ura4* in pDB4697. *Padh1-OsTIR1-F74A* integrating plasmid pDB4695 was constructed by mutating the codon for amino acid 74 of OsTIR1 from TTT (Phe) to GCT (Ala). Primers used for plasmid construction are listed in Table S1. pDB4581, pDB4697, pDB5050, and pDB4695 have been deposited at Addgene (Addgene IDs 171124, 171125, 171126, and 171127, respectively).

### Strain construction

Fission yeast strains used in this study are listed in Table S2. Strains constructed by Kanke et al. were acquired from the Yeast Genetic Resource Center of Japan (YGRC/NBRP) (http://yeast.nig.ac.jp/). The IAA17-tagged alleles and the F-box-protein-expressing cassettes generated by Kanke et al. were combined with AID components created in this study by crossing. To integrate the *Padh1-OsTIR1* cassette or the *Padh1-OsTIR1-F74A* cassette into the genome, we transformed a *ura4-D18* strain with NotI-digested pDB5050 or pDB4695. Ends-out recombination led to the integration of the cassette and converted the strain to *ura4^+^*. Integration at the *ura4* locus was verified by PCR. Endogenous C-terminal tagging of target genes with the 3×sAID degron was performed using a PCR-based tagging method. Briefly, in step 1, the degron cassette was amplified from pDB4581 using primers TAG-F and TAG-R and two homology arms were amplified from genomic DNA using the primer pair of orf_F and up_ov_R3 and the primer pair of dn_ov_F3 and dn _R, respectively. In step 2, overlap PCR using the three PCR products from step 1 as templates and orf_F and dn_R as primers generated the final PCR product used for transformation. Transformation mix was spread on YES plates, incubated at 30°C for 2 days, and then replica plated on G418-containing YES plates. Primers used for strain construction are listed in Table S1.

### Chemical synthesis of 5-adamantyl-IAA

5-adamantyl-IAA was synthesized using a protocol modified from two previously reported procedures (Samann et al. 2014; Petit et al. 2009). First, 5-adamantyl-indole was generated through Negishi coupling of 5-iodoindole and an adamantylzinc reagent, which was prepared from the corresponding adamantylmagnesium reagent via transmetallation. 5-adamantyl-IAA was then synthesized from 5-adamantyl-indole via glyoxylation and a selective ketone reduction procedure. Full experimental details were presented in Fig. S1.

We note that 5-adamantyl-IAA has recently become available from several commercial vendors including Tokyo Chemical Industry (product number A3390).

### Culturing media

Cells were cultured at 30°C in pombe minimal medium with glutamate (PMG) containing necessary supplements. The composition of this medium was as described (Forsburg and Rhind 2006). For preparing NAA-containing media, we added into the medium 0.5 M NAA stock solution (dissolved in 1 M NaOH and stored at -20°C) and, if necessary, adjusted the pH of the medium to 6.0 using 1 M HCl. For preparing 5-adamantyl-IAA-containing media, we added into the medium 1 mM 5-adamantyl-IAA stock solution (dissolved in 1 M NaOH and stored at -20°C) and, if necessary, adjusted the pH of the medium to 6.0 using 1 M HCl. The stock solutions were diluted in water before an appropriate amount was added to the medium. The pH of the medium was not obviously affected by NaOH in the stock solution and no adjustment of pH was needed if the final dilution ratio was 1000 folds or higher, i.e. if the final concentration of NAA ≤ 0.5 mM and if the final concentration of 5-adamantyl-IAA ≤ 1 μM. NAA was from Sigma (catalog number N0640).

### Spot assay

For the spot assay, cultures were grown to log phase in a liquid medium and serial 5-fold dilutions of cells were spotted onto PMG plates, which were incubated at 30°C and scanned using an Epson Perfection V800 photo scanner after colonies formed on the plates.

### Immunoblotting

Cell lysates were prepared using a trichloroacetic acid extraction method (Ulrich and Davies 2009). Samples were separated on a 10% SDS-PAGE and immunoblotted with an anti-mini-AID antibody (MBL, Code No. M214-3).

### RNA-seq analysis

A log-phase culture of strain DY38751 grown in PMG liquid medium was diluted to OD600 = 0.2 and split into six equal aliquots. Into two aliquots, we added 2 mM NAA (250-fold dilution from 0.5 M NAA stock dissolved in 1 M NaOH). Into another two aliquots, we added 4 µM 5-adamantyl-IAA (250-fold dilution of 1 mM 5-adamantyl-IAA stock dissolved in 1 M NaOH). The remaining two aliquots were added with equivalent amounts of 1 M NaOH and served as controls. 1 M HCl was added to the six aliquots to adjust the pH to 6.0. These six cultures were incubated at 30°C for 6 h. Cells were then collected and RNA was isolated using the hot phenol method (Lyne et al. 2003). RNA-seq library preparation and Illumina sequencing were performed by Annoroad Gene Technology (Beijing, China).

The RNA-seq sequencing reads were aligned to the *S. pombe* reference genome sequence using HISAT2 version 2.1.0 with the options ’--min-intronlen 28 --max-intronlen 820’ (Kim et al. 2019). The CDS coordinates of protein-coding genes were from the annotation file Schizosaccharomyces_pombe.ASM294v2.44.gtf (ftp://ftp.ensemblgenomes.org/pub/fungi/release-44/gtf/schizosaccharomyces_pombe/). Reads mapped to each CDS were counted by featureCounts version 2.0.1 with the options ’-p -s 0 -F SAF’ (Liao et al. 2014). Differential expression analysis was performed using the R package DESeq2 version 1.28.1 (Love et al. 2014). Volcano plots were generated using the R package EnhancedVolcano version 1.6.0 (https://github.com/kevinblighe/EnhancedVolcano). Sequencing data have been deposited at NCBI SRA under the BioProject ID PRJNA743700. The accession numbers of the data of the control samples are SRR15041253 and SRR15041254. The accession numbers of the data of the 5-adamantyl-IAA-treated samples are SRR15041255 and SRR15041256. The accession numbers of the data of the NAA-treated samples are SRR15041257 and SRR15041258.

### Data Availability Statement

Strains and plasmids are available upon request. The authors affirm that all data necessary for confirming the conclusions of the article are present within the article, figures, and tables.

## RESULTS

### *Padh1* promoter-expressed OsTIR1 is more effective than Skp1-fused F-box proteins used in the Kanke system

It is known that in AID systems, high expression levels of the plant F-box proteins are crucial for efficient protein depletion (Nishimura et al. 2009; Kanke et al. 2011; Nishimura and Kanemaki 2014; Zhang et al. 2015; Natsume et al. 2016; Natsume and Kanemaki 2017; Texier et al. 2018). In the Kanke system, the F-box protein AtTIR1 is fused with *S. pombe* Skp1 and two copies of SV40 nuclear localization signal (NLS) and is expressed from the moderate-strength *Padh15* promoter (Kanke et al. 2011). Because in many other species, plant F-box proteins have been successfully used without fusing to endogenous Skp1 (Nishimura et al. 2009; Kanke et al. 2011; Nishimura and Kanemaki 2014; Zhang et al. 2015; Natsume et al. 2016; Natsume and Kanemaki 2017; Texier et al. 2018), we envisioned that Skp1 fusion may not be necessary for *S. pombe*. Expressing the F-box proteins without Skp1 fusion may avoid toxicity and allow the F-box proteins to be expressed from a strong promoter. Thus, we decided to not fuse the F-box protein with Skp1 in our system.

The choice of fusing AtTIR1 with two copies of NLS in the Kanke system stemmed from the intention of only applying the system to degrade nuclear proteins. When AtTIR1 is concentrated inside the nucleus by the NLSs, it may no longer be effective at degrading cytoplasmic proteins. We aimed to develop a system that can be used for degrading both nuclear and cytoplasmic proteins. Because F-box proteins are relatively small (less than 70 kDa), they do not need an NLS to enter the nucleus and in the absence of NLS-dependent active transport into the nucleus, they can still passively diffuse through nuclear pores on the timescale of minutes (Timney et al. 2016). Therefore, we chose to not add an NLS to the F-box protein in our system.

An OsTIR1 coding sequence codon-optimized for *S. pombe* was obtained by gene synthesis and placed under the control of the strong *Padh1* promoter (Fig. 1A). The OsTIR1-expressing cassette (hereafter referred to as *Padh1-OsTIR1*) was integrated at the *ura4* locus by ends-out recombination using the plasmid pDB5050 (Table 1 and Fig. S2A). Codon-optimized OsTIR1 expressed from the strong *Padh1* promoter showed no obvious toxicity in the absence of auxin and the AID degron (Fig. S2B).

**Figure 1.**
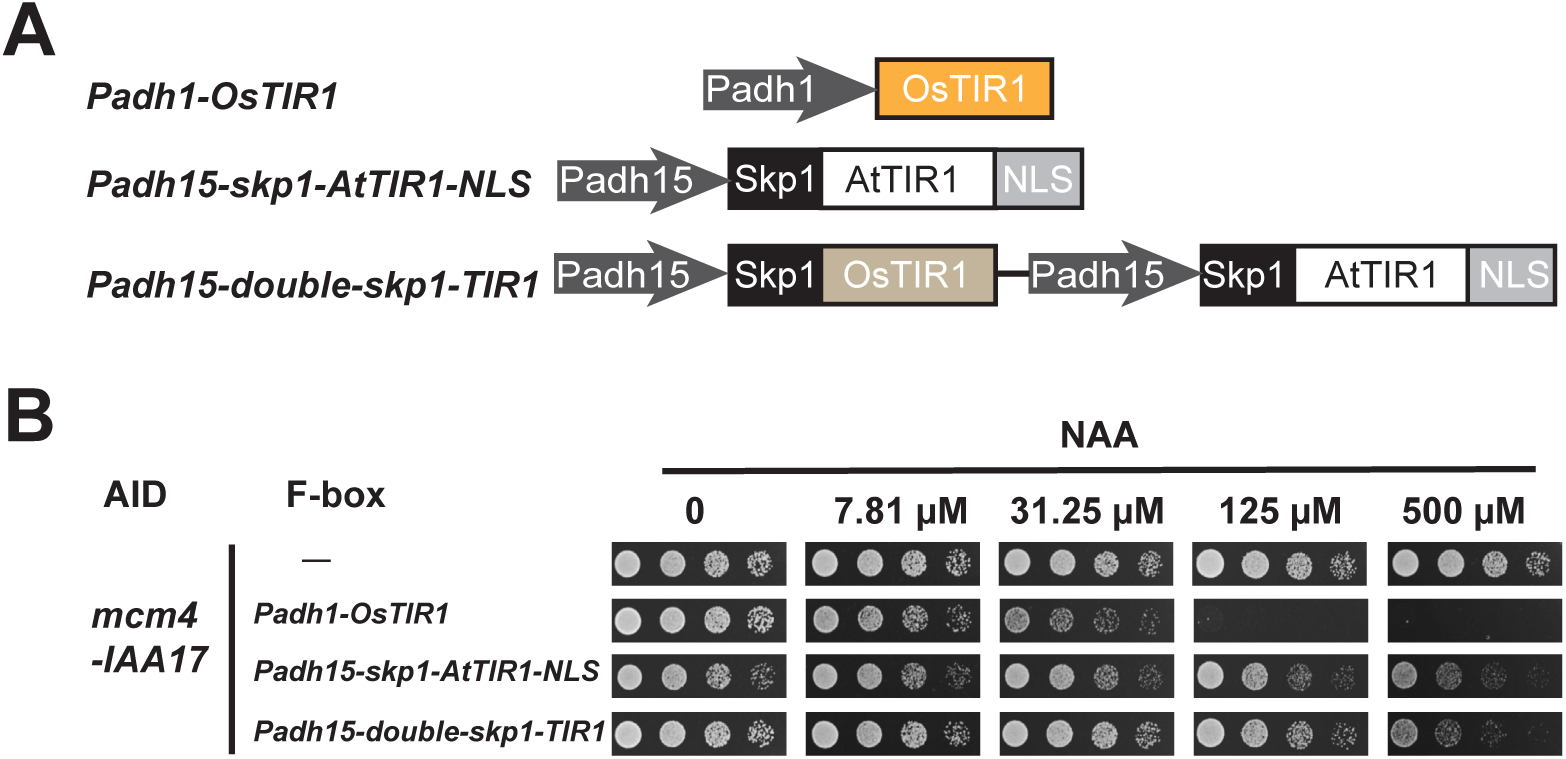
*Padh1* promoter-expressed OsTIR1 is more effective than F-box proteins used in the Kanke system. A. Schematic of the genome-integrated cassettes expressing F-box proteins (not drawn to scale). The *Padh1-OsTIR1* cassette expressing codon-optimized OsTIR1 from the strong *Padh1* promoter was designed and constructed in this study. The *Padh15-skp1-AtTIR1-NLS* cassette and the *Padh15-double-skp1-TIR1* cassette were developed by Kanke et al. (Kanke et al. 2011). B. Spot assay showed that growth inhibition occurred at a lower concentration of NAA when the *mcm4-IAA17* allele was combined with the *Padh1-OsTIR1* cassette than when combined with the *Padh15-skp1-AtTIR1-NLS* cassette or the *Padh15-double-skp1-TIR1* cassette.

To test the effectiveness of the *Padh1* promoter-expressed OsTIR1 in an AID system, we targeted the essential gene *mcm4*, which encodes a nuclear protein required for DNA replication. For essential genes, effective protein depletion manifests as growth retardation. We used the degron-tagged *mcm4* allele generated by Kanke et al., *mcm4-IAA17*, in which the full-length IAA17 was used as the degron. Kanke et al. previously developed two versions of integrated F-box-protein-expressing cassettes, *Padh15-skp1-AtTIR1-NLS* and *Padh15-double-skp1-TIR1* (Fig. 1A). Both versions, when combined with the *mcm4-IAA17* allele, resulted in obvious but incomplete growth inhibition at 30°C when the inducer NAA was added at the concentration of 500 µM, which is the concentration previously used by Kanke et al. (Fig. 1B). In contrast, when our *Padh1-OsTIR1* cassette was combined with the *mcm4-IAA17* allele, complete growth inhibition was observed at 125 µM of NAA (Fig. 1B). Because *Padh1-OsTIR1* resulted in a stronger growth inhibition at a 4-fold lower concentration of inducer, we concluded that *Padh1-OsTIR1* is more effective than the F-box-protein-expressing cassettes of the Kanke system for disrupting the function of *mcm4*.

To determine whether the higher effectiveness of the *Padh1-OsTIR1* cassette holds true in general, we targeted two other essential genes *cdc45* and *orc2* (Fig. S3). Using the *cdc45-IAA17* and *orc2-IAA17* alleles generated by Kanke et al., we found that for both genes, *Padh1-OsTIR1* was more potent than the F-box-protein-expressing cassettes developed by Kanke et al. (Fig. S3). Together, these results indicate that *Padh1-OsTIR1* is an improvement of the AID system for *S. pombe*.

### 3×sAID is a more effective degron than the full-length IAA17

In the Kanke system, the full-length IAA17 is used as the degron (Kanke et al. 2011). To reduce the size of the degron and thereby minimize the possibility of the fused degron interfering with the function of the target protein, we selected a 36-amino-acid sequence from the AtIAA17 protein (amino acids 71-106 of AtIAA17, hereafter referred to as sAID, for short AID) based on sequence conservation and known structure-function relationship of Aux/IAA proteins (Fig. 2A). sAID is shorter than previously reported small-size AID degrons derived from IAA17, including AID*, AID^47^, and mini-AID (Kubota et al. 2013; Morawska and Ulrich 2013; Brosh et al. 2016; Natsume et al. 2016; Yesbolatova et al. 2019) (Fig. S4A). Based on a previous report showing that three copies of mini-AID worked better than a single copy of mini-AID (Kubota et al. 2013), we used three tandem copies of the sAID sequence as the degron (hereafter referred to as 3×sAID) (Figure 2A). To avoid recombination between the three copies, we introduced synonymous variations in the three copies, so that between-copy nucleotide identities are lower than 80% (Fig. S4B).

**Figure 2.**
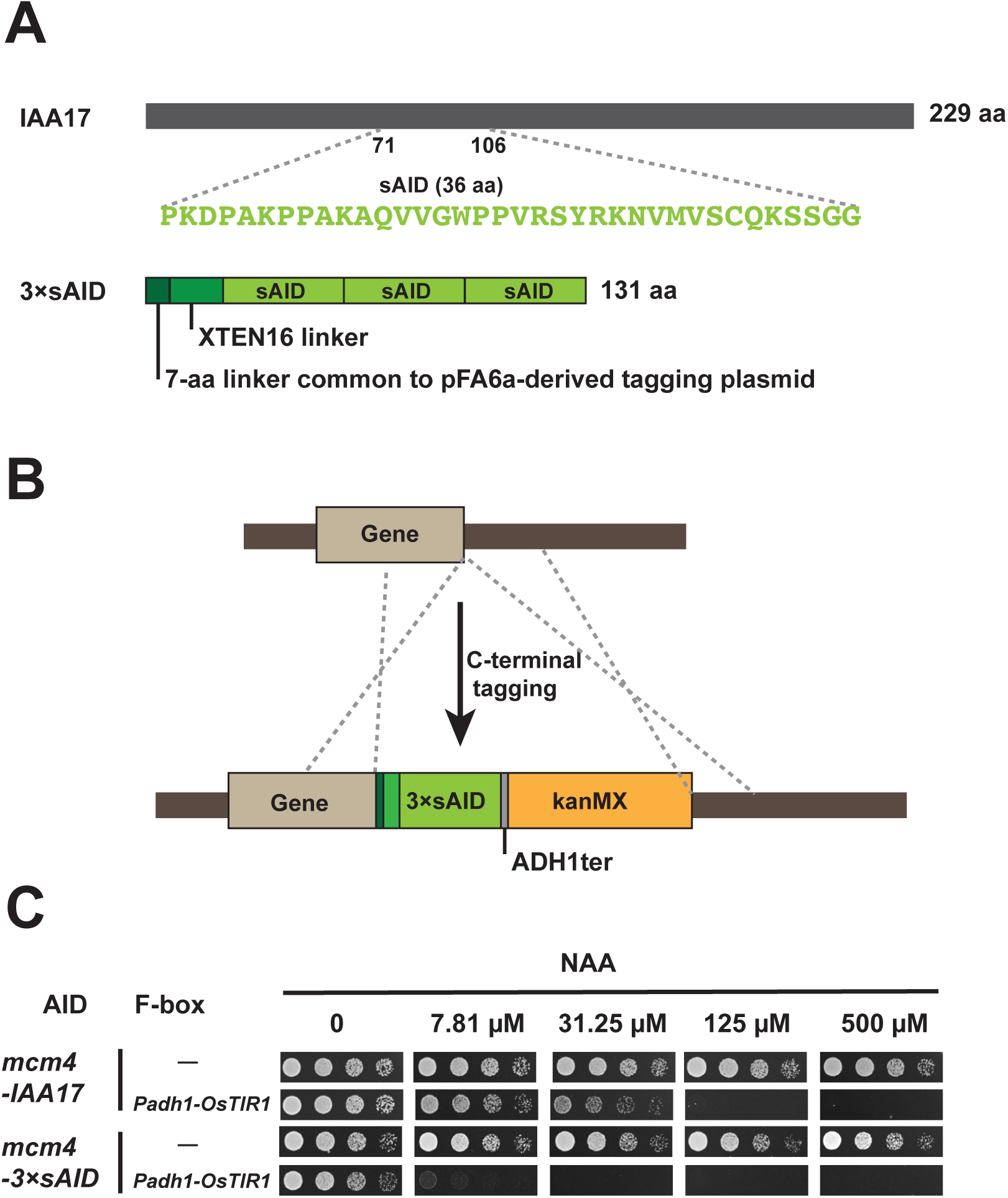
3×sAID is a more effective degron than the full-length IAA17. A. Schematic of the 3×sAID degron in the tagging plasmid pDB4581. B. Schematic of endogenous C-terminal tagging of a gene with the 3×sAID degron in the plasmid pDB4581 (not drawn to scale). *ADH1ter* and *kanMX* are the terminator of *S. cerevisiae ADH1* gene and the G418-resistance selection marker, respectively. C. Spot assay showed that growth inhibition occurred at a lower concentration of NAA when the *Padh1-OsTIR1* cassette was combined with *mcm4-3×sAID* than when combined with *mcm4-IAA17*.

We constructed a C-terminal 3×sAID tagging plasmid (pDB4581) (Table 1 and Fig. 2B), which is based on the pFA6a-derived tagging plasmid series (Bahler et al. 1998; Longtine et al. 1998). In this plasmid, upstream of the 3×sAID degron, we added a 16-amino-acid XTEN16 linker (SGSETPGTSESATPES), which was initially developed by David Liu’s lab for linking Cas9 and FokI and has been adopted by Jonathan Weissman’s lab as a linker between an AID tag and the target protein (Guilinger et al. 2014; Costa et al. 2018). Like other pFA6a-derived C-terminal tagging plasmids, when using pDB4581 as a template for PCR-based C-terminal tagging, a 7-amino-acid linker (RIPGLIN) is present immediately downstream of the target protein. The total length of the amino acid sequence added to the target protein, including the 3×sAID degron and the linkers, is only 131 amino acids, substantially shorter than the length of full-length IAA17 (229 amino acids).

To test whether the 3×sAID degron is more effective than the full-length IAA17 used in the Kanke system, we endogenously tagged Mcm4 with the 3×sAID degron. When *mcm4-3×sAID* was combined with *Padh1-OsTIR1*, nearly complete growth inhibition was observed at 7.81 µM of NAA (Fig. 2C). Because *mcm4-IAA17* in combination with *Padh1-OsTIR1* did not show as severe a growth phenotype at a 4-fold higher concentration of NAA (31.25 µM) (Fig. 2C), we concluded that 3×sAID is a more effective degron than the full-length IAA17 for targeting *mcm4*.

We tested the new degron on two additional essential genes *pol1* and *cdc20* (Fig. S4). In combination with *Padh1-OsTIR1*, *pol1-IAA17* allele previously constructed by Kanke et al. exhibited a moderate growth inhibition at 500 µM of NAA, whereas *pol1-3×sAID* allele exhibited a nearly complete growth inhibition at 31.25 µM of NAA (Fig. S5A). For *cdc20*, in combination with *Padh1-OsTIR1*, the *cdc20-IAA17* allele previously constructed by Kanke et al. showed no obvious growth phenotype at 500 µM of NAA, whereas the same concentration of NAA completely inhibited the growth of *pol1-3×sAID* cells (Fig. S5B). Together, these results indicate that 3×sAID is superior to the full-length IAA as a degron in AID systems.

### 5-adamantyl-IAA, when paired with OsTIR1-F74A, is a highly effective inducer

Because auxins including IAA and NAA are known to have physiological effects on yeasts, such as growth impediment and perturbation of TORC1, we decided to adopt a synthetic auxin analog, 5-adamantyl-IAA, as the inducer for the AID system. Compared to IAA, 5-adamantyl-IAA bears an extra bulky adamantane group (Fig. 3A), and thus is unlikely to cause the same physiological effects as auxins cause. Furthermore, 5-adamantyl-IAA can induce an interaction between the F79A mutant form of AtTIR1, which has an enlarged auxin-binding pocket, and Aux/IAA proteins at an extremely low concentration of 10 pM (Uchida et al. 2018; Torii et al. 2018; Yamada et al. 2018), suggesting the possibility of using 5-adamantyl-IAA as an AID inducer at much lower concentrations than auxins. In OsTIR1, the F74A mutation is equivalent to the F79A mutation in AtTIR1. We introduced the F74A mutation into codon-optimized OsTIR1 of the *Padh1-OsTIR1* cassette and generated a plasmid for integrating the *Padh1-OsTIR1-F74A* cassette at the *ura4* locus through ends-out recombination (pDB4695) (Fig. 3B and Table 1). Like OsTIR1 expressed from the *Padh1* promoter, OsTIR1-F74A expressed from the *Padh1* promoter did not cause obvious growth inhibition in the absence of auxin and the AID degron (Fig. S2B).

**Figure 3.**
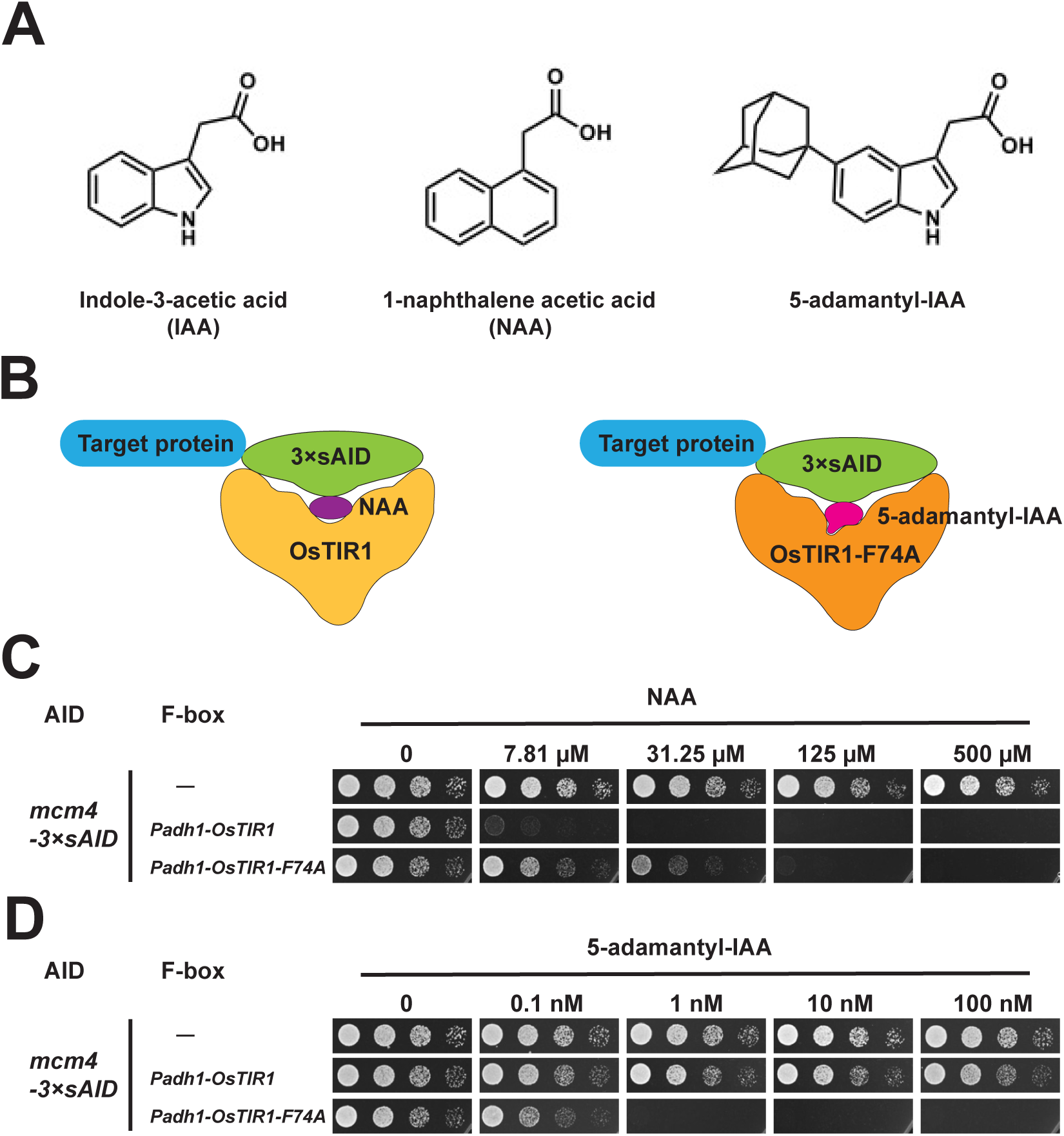
5-adamantyl-IAA is a highly effective inducer when paired with OsTIR1-F74A. A. The structures of IAA, NAA, and 5-adamantyl-IAA. B. Schematic showing that NAA preferentially binds OsTIR1, whereas 5-adamantyl-IAA preferentially binds OsTIR1-F74A. C. Spot assay showed that *Padh1-OsTIR1-F74A* was less effective than *Padh1-OsTIR1* when using NAA as the inducer. D. Spot assay showed that *Padh1-OsTIR1-F74A* but not *Padh1-OsTIR1* was effective when using 5-adamantyl-IAA as the inducer. Spot assay was performed using PMG plates incubated at 30°C.

We compared the abilities of *Padh1-OsTIR1* and *Padh1-OsTIR1-F74A* to generate growth inhibitory effects on cells harboring the *mcm4-3×sAID* allele in response to the two different inducers NAA and 5-adamantyl-IAA (Fig. 3C and Fig. 3D). As expected, *Padh1-OsTIR1-F74A* was less effective than *Padh1-OsTIR1* when using NAA as the inducer (Fig. 3C), presumably because the F74A mutation makes NAA not fit well in the auxin-binding pocket. When using 5-adamantyl-IAA as the inducer, we found that, remarkably, 1 nM of 5-adamantyl-IAA was enough to completely inhibit the growth of cells containing the *Padh1-OsTIR1-F74A* cassette (Fig. 3D). In contrast, hardly any growth inhibitory effect was observed at 100 nM of 5-adamantyl-IAA for cells containing the *Padh1-OsTIR1* cassette, consistent with the expectation that the bulky adamantane group of 5-adamantyl-IAA makes it unable to fit into the auxin-binding pocket of wild-type OsTIR1 (Fig. 3D). The fact that 1 nM of 5-adamantyl-IAA in combination with *Padh1-OsTIR1-F74A* generated a stronger growth inhibition than 7.81 µM of NAA in combination with *Padh1-OsTIR1* (comparing the second row in Fig. 3C and the third row in Fig. 3D) suggests that 5-adamantyl-IAA is a highly effective inducer when paired with OsTIR1-F74A and can be used at concentrations thousands of times lower than NAA used in combination with wild-type OsTIR1. We concluded that the combination of the 3×sAID degron, the *Padh1-OsTIR1-F74A* cassette, and 5-adamantyl-IAA as the inducer constitute an improved AID system for *S. pombe* (Fig. 4).

**Figure 4.**
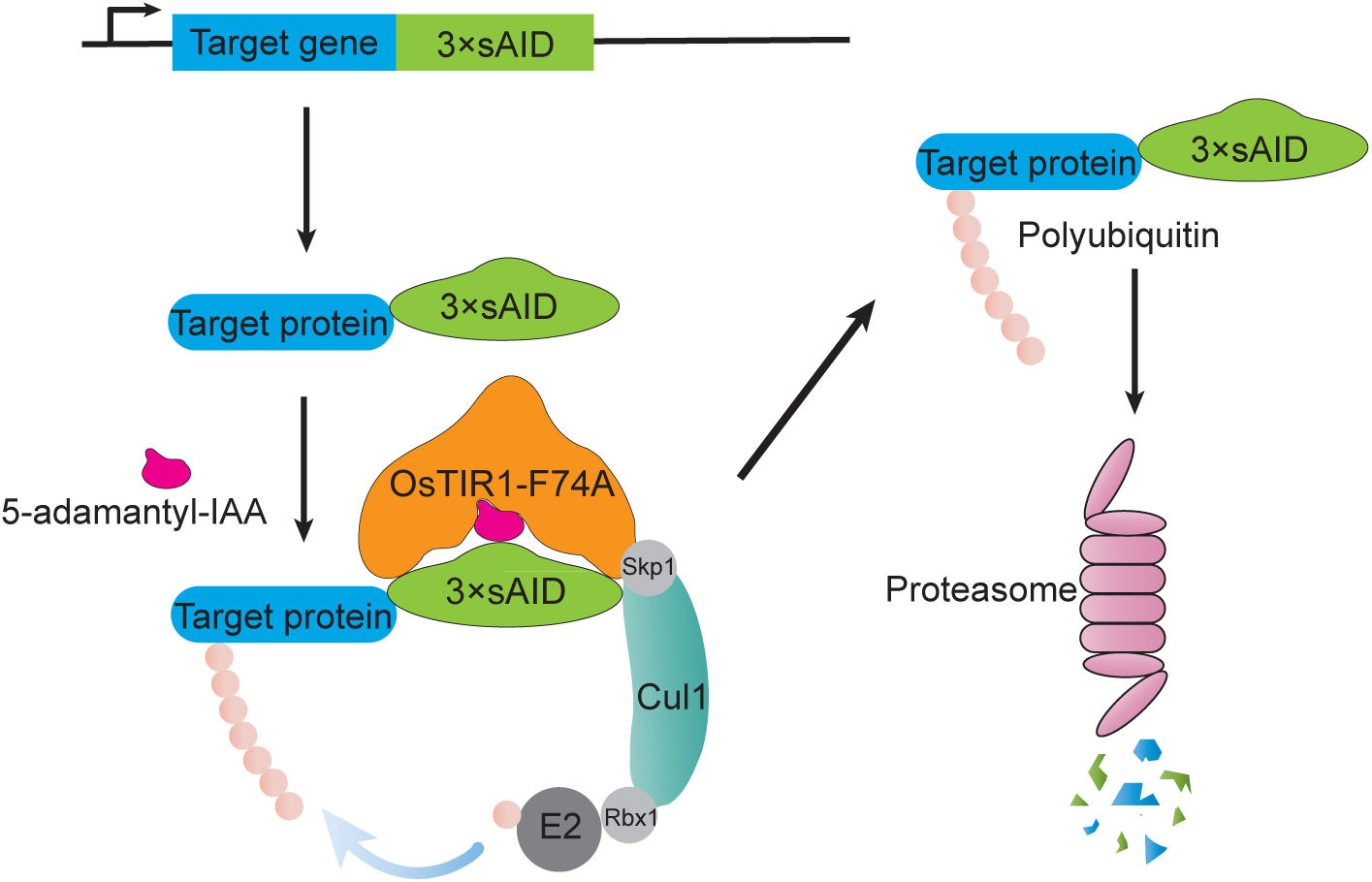
The improved AID system for *S. pombe*. Schematic of the improved AID system. OsTIR1-F74A can form a functional E3 ligase complex with endogenous SCF components. 5-adamantyl-IAA acts as a molecule glue to promote the interaction between OsTIR1-F74A and 3×sAID, resulting in the polyubiquitination and degradation of the 3×sAID fused target protein.

To directly assess the extent and rapidity of protein depletion driven by our improved AID system, we performed immunoblotting analysis (Fig. S6). Complete depletion of Mcm4-3×sAID was observed upon treating the cells with 5-adamantyl-IAA for 15 min, indicating that rapid and complete protein depletion can be achieved using the improved AID system.

### Our improved AID system is effective on genes refractory to the Kanke system

Kanke et al. has tested their AID system on 15 essential genes, all of which encode nuclear proteins, and found that for only 3 of these 15 genes, severe growth phenotype can be obtained (Kanke et al. 2011). We applied our improved AID system on 8 genes previously tested by Kanke et al. (Table 2), including 2 genes (*mcm4* and *orc2*) that exhibited severe growth defect using the Kanke system, 2 genes (*cdc45* and *orc6*) that exhibited moderate growth defect using the Kanke system, and 4 genes (*pol1*, *cdc20*, *mcm10*, and *ssl3*) that exhibited no growth phenotype using the Kanke system. The improved AID system was able to generate induced lethality (complete growth inhibition on a medium containing 5-adamantyl-IAA) for all 8 genes encoding nuclear proteins. Consistent with the results of Kanke et al., higher concentrations of 5-adamantyl-IAA was required to generate induced lethality for the genes that exhibited weaker phenotype in the Kanke system (Table 2). *cdc20*, *mcm10*, and *ssl3* required the highest concentration of 5-adamantyl-IAA (100 nM) for complete growth inhibition. We concluded that our improved AID system permits inactivation of genes refractory to the Kanke system and 100 nM of 5-adamantyl-IAA should be sufficient for generating a strong inactivating effect for most target genes.

**Table 2.** Our improved AID system is effective on genes that are refractory to the Kanke system.

| Essential gene | Results obtained using the Kanke system (according to Kanke et al.) | Results obtained using our improved AID system (the concentration of 5-adamantyl-IAA that resulted in complete growth inhibition) * |
| --- | --- | --- |
| *mcm4* | Severe growth defect | Complete growth inhibition at 1 nM |
| *orc2* | Severe growth defect | Complete growth inhibition at 10 nM |
| *cdc45* | Slow growth | Complete growth inhibition at 10 nM |
| *orc6* | Slow growth | Complete growth inhibition at 10 nM |
| *pol1* | No effect | Complete growth inhibition at 10 nM |
| *cdc20* | No effect | Complete growth inhibition at 100 nM |
| *mcm10* | No effect | Complete growth inhibition at 100 nM |
| *ssl3* | No effect | Complete growth inhibition at 100 nM |

\*Growth phenotype was examined by spot assays using PMG plates containing 4 different concentrations of 5-adamantyl-IAA (0.1, 1, 10, and 100 nM) at 30°C.

### The improved AID system can be used for inactivating genes encoding cytoplasmic proteins

Kanke et al. has only tested their system on genes encoding nuclear proteins (Kanke et al. 2011). In our system, we intentionally opted to not add an NLS to the F-box protein OsTIR1 so that it can target both nuclear and cytoplasmic proteins. To test whether the improved AID system is effective in disrupting the functions of genes encoding cytoplasmic proteins, we endogenously tagged two cytoplasmic proteins, Sec8 and Sec16, with 3×sAID at their C-termini. Both Sec8 and Sec16 are essential for growth. Complete growth inhibition was observed with 100 nM of 5-adamantyl-IAA for both *sec8*-3*×sAID* and *sec16*-3*×sAID* (Fig. 5). These results suggest that our improved AID system can efficiently deplete cytoplasmic proteins in *S. pombe*.

**Figure 5.**
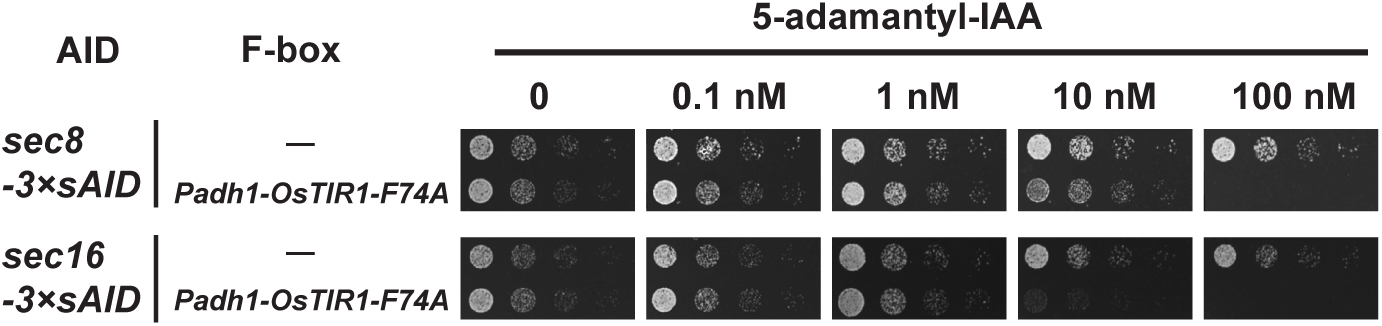
The improved AID system is effective on cytoplasmic proteins. Spot assay showed that complete growth inhibition was achieved when using the improved AID system to disrupt the functions of two essential genes encoding cytoplasmic proteins.

### NAA but not 5-adamantyl-IAA perturbs the transcriptome

To experimentally assess whether NAA and 5-adamantyl-IAA may cause undesirable side effects in *S. pombe*, we performed RNA-seq analysis on *S. pombe* cells containing neither an AID degron nor a TIR1 protein. We treated cells with either 2 mM of NAA or 4 μM of 5-adamantyl-IAA for 6 h. 2 mM of NAA is 4 times the concentration recommended by Kanke et al. for the Kanke system (Kanke et al. 2011); 4 μM of 5-adamantyl-IAA is 40 times the maximal concentration we used with our improved AID system. In our hands, neither 2 mM of NAA nor 4 μM of 5-adamantyl-IAA caused an obvious growth phenotype (our unpublished observations). Using fold changes > 2 and adjusted *P* values < 0.05 as cutoffs for differentially expressed genes, we found that 18 genes were up-regulated and 10 genes were down-regulated upon NAA treatment (Fig. 6 and Table S3). In contrast, using the same cutoffs, no differentially expressed genes were found in cells treated with 5-adamantyl-IAA (Fig. 6 and Table S3). These results indicate that a high concentration of NAA can alter the transcriptome of *S. pombe*, whereas 5-adamantyl-IAA at a concentration far higher than those used in our improved AID system had no obvious effect on the transcriptome.

**Figure 6.**
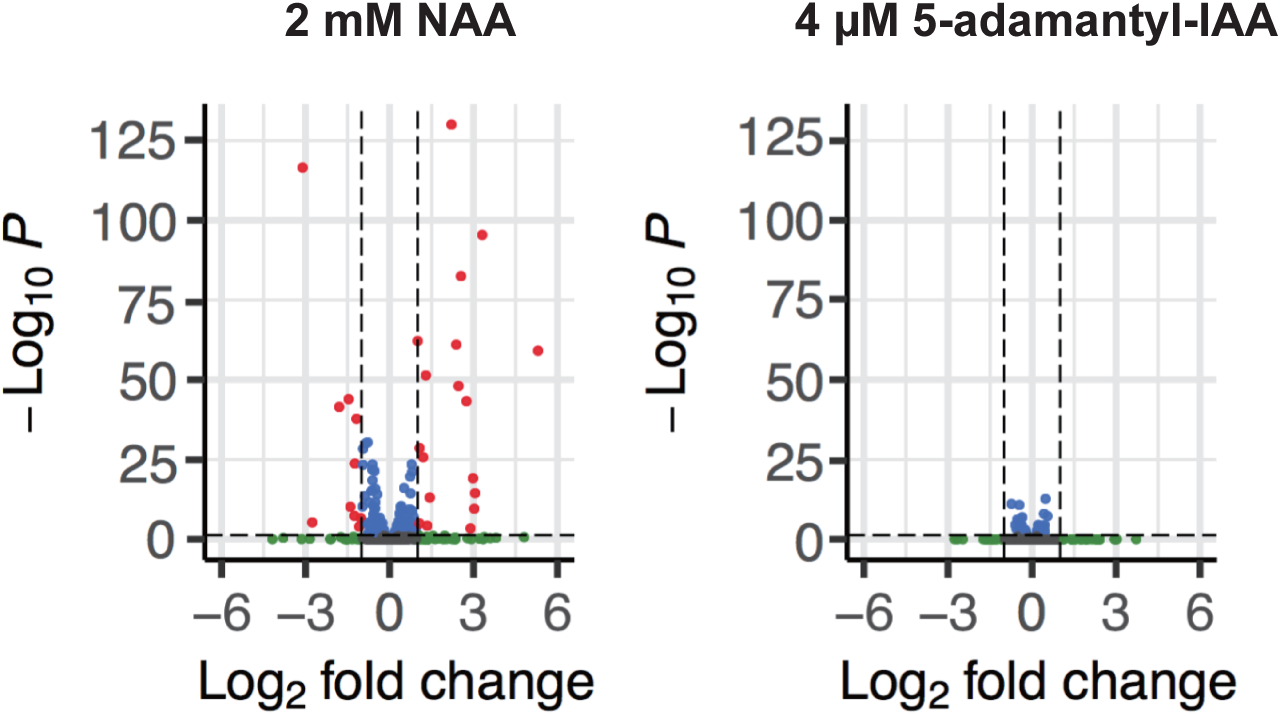
2 mM of NAA but not 4 μM of 5-adamantyl-IAA perturbed the transcriptome. Changes in the transcriptome are visualized using volcano plots. Left, NAA-treated samples are compared with control samples; right, 5-adamantyl-IAA-treated samples are compared with control samples. The x-axis shows the Log2 (fold change) in transcript levels between treated and control samples. The two vertical dashed lines denote Log2 (fold change) = −1 and 1, respectively. The y-axis shows the −Log10 (adjusted *P* value). The horizontal dashed line denotes adjusted *P* value = 0.05. Genes with fold changes > 2 but adjusted *P* values > 0.05 are shown in green, genes with adjusted *P* values < 0.05 but fold changes < 2 are shown in blue, and genes with fold changes > 2 and adjusted *P* values < 0.05 are shown in red. The smallest adjusted *P* value for NAA treatment, that of the gene SPCC569.05c, is 5.5654E-197. For better visualization, we arbitrarily set the adjusted *P* value of SPCC569.05c to 1E-130. For genes that were given NA values for adjusted *P* values by DESeq2, we set their adjusted *P* values to 1.

## DISCUSSION

Here, we report an improved AID system that permits rapid and efficient protein depletion in fission yeast. In this system, we combine an improved degron composed of three tandem copies of a 36-amino-acid sequence from AtIAA17, codon-optimized OsTIR1-F74A expressed from a strong promoter, and 5-adamantyl-IAA as an inducer to minimize side effects. This system is effective on disrupting the functions of 10 essential genes, including genes that are refractory to a previously reported system.

During the preparation of our manuscript, another study on improving the AID system for *S. pombe* was published (Watson et al. 2021). Similar to our findings, Watson et al. also showed that OsTIR1 not fused with Skp1 can be expressed from a strong promoter without causing toxicity and is more effective in inactivating *mcm4* than the *Padh15-skp1-AtTIR1-NLS* cassette of the Kanke system. Like the Kanke system, Watson et al. aimed to use their system for depleting nuclear proteins and therefore added an NLS to OsTIR1. They showed that two nuclear proteins, Mcm4 and Rad52, were effectively depleted. It is likely that using an NLS to target OsTIR1 to the nucleus can lower the efficiency of depleting cytoplasmic proteins. In our system, we intentionally chose to not add an NLS to OsTIR1 to allow broad applicability. Indeed, genes encoding both nuclear and cytoplasmic proteins can be effectively inactivated using our improved system.

We showed in this study that, when paired with OsTIR1-F74A, 5-adamantyl-IAA is a highly effective inducer for the AID system and can minimize side effects on the transcriptome. Several recently published studies have also demonstrated that a bulky auxin analog, when paired with a mutant form of OsTIR1 with an enlarged auxin-binding pocket, acts as an effective AID inducer at much lower concentrations than auxin (Yesbolatova et al. 2020; Nishimura et al. 2020; Watson et al. 2021). When we initiated this study in 2018, 5-adamantyl-IAA was not commercially available and therefore we obtained it by chemical synthesis. Recently, it has become available from several commercial vendors. Because 5-adamantyl-IAA can be used at very low concentrations, it is a highly cost-effective inducer. For example, the amount of 5-adamantyl-IAA needed to prepare 1 liter of medium containing 100 nM of 5-adamantyl-IAA costs less than half a US dollar.

In mammalian cell lines, OsTIR1-based AID systems have been reported to cause the degradation of degron-fused target proteins and generate loss-of-function phenotypes in the absence of the auxin inducer (Natsume et al. 2016; Yesbolatova et al. 2019; Li et al. 2019; Sathyan et al. 2019). In budding yeast, auxin-independent depletion of AID-tagged proteins was also observed when OsTIR1 was expressed to a high level (Mendoza-Ochoa et al. 2019). This undesirable phenomenon, called “basal degradation” or “leaky degradation”, has been tackled by a number of approaches, including placing OsTIR1 under the control of a repressible promoter (Natsume et al. 2016; Mendoza-Ochoa et al. 2019), adding a small molecule antagonist of OsTIR1 (Yesbolatova et al. 2019), switching to a different F-box protein (Li et al. 2019), expressing another plant auxin signaling protein ARF (Sathyan et al. 2019), and using the OsTIR1-F74G mutant (Yesbolatova et al. 2020). It was reported that in budding yeast, OsTIR1-F74A, when expressed from the *GAL1-10* promoter, can cause inducer-independent loss-of-function phenotypes (Yesbolatova et al. 2020). In fission yeast, we have not noticed any obvious inducer-independent phenotypes using either wild-type OsTIR1 or OsTIR1-F74A expressed from the strong *Padh1* promoter (spot assay data shown in this paper and our unpublished observations). Furthermore, our immunoblotting analysis showed that, in the absence of inducer, OsTIR1-F74A expressed from the strong *Padh1* promoter did not obviously alter the protein level of Mcm4-3×sAID (Fig. S6). Thus, under the experimental conditions we have used, basal degradation is not an apparent problem with our improved AID system for fission yeast. If this problem arises under other experimental conditions, the myriad of solutions mentioned above can be applied to address it.

The strengths and advantages of *S. pombe* as an experimental model organism rely on the continuing development of technologies empowering its use in biological research. Understanding gene functions is a core pursuit of biological research. Our improved AID system is a new addition to a long list of tools and methods for creating conditional loss-of-function mutants in *S. pombe* (Basi et al. 1993; Rajagopalan et al. 2004; Boe et al. 2008; Kearsey and Gregan 2009; Kanke et al. 2011; Tang et al. 2011; Pai et al. 2012; Watson et al. 2013; Ding et al. 2014; Watson et al. 2021). This system will facilitate the analysis of gene functions in *S. pombe*, especially the functions of essential genes.

## ACKNOWLEDGMENTS

We thank Hisao Masukata for making the strains described in Kanke et al. available through YGRC/NBRP, and YGRC/NBRP for providing the strains. This work was supported by grants from the Ministry of Science and Technology of the People’s Republic of China and the Beijing municipal government.

## Supplementary Figures

**Figure S1.**
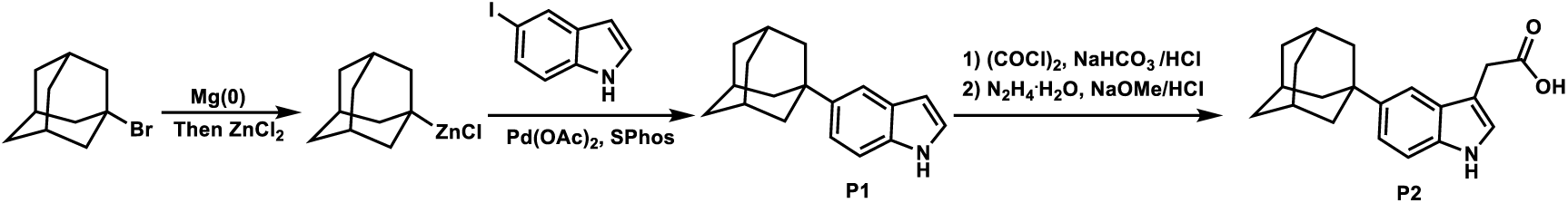
Schematic of the synthesis of 5-adamantyl-IAA (P2). The procedure of **P1** synthesis: A dry, argon flushed Schlenk-flask equipped with a magnetic stirring bar and a septum was charged with LiCl (187 mg, 4.4 mmol, 1.1 equiv) and heated with a heat gun under a high vacuum (5 min). After cooling to room temperature, magnesium turnings (194 mg, 8.0 mmol, 2 equiv) were added, followed by THF (10 mL). The magnesium was activated using 1,2-dibromoethane (38 mg, 0.2 mmol, 5 mol%) and TMSCl (22 mg, 0.2 mmol, 5 mol%). After cooling to 0°C, ZnCl_2_ (600 mg, 4.4 mmol, 1.1 equiv) was added followed by the 1-bromoadamantane (861 mg, 4.0 mmol, 1 equiv). The reaction mixture was stirred at 25°C for 5 h. Then 5-iodo-1*H*-indole (875 mg, 3.6 mmol, 0.9 equiv) was added to the freshly prepared zinc reagent followed by Pd(OAc)_2_ (9 mg, 0.04 mmol, 1 mol%) and SPhos (33 mg, 0.08 mmol, 2 mol%) and the mixture was stirred at 50°C for 5 h. The reaction mixture was quenched with sat. NH_4_Cl solution (10 mL) and extracted with EtOAc (3×20 mL). The combined organic phases were dried over Na_2_SO4 and concentrated *in vacuo*. The crude residue obtained was purified by flash column chromatography to give the pure product **P1** (white solid, 610 mg, 60% yield). ^1^H NMR (400 MHz, Chloroform-*d*) δ 8.05 (br s, 1H), 7.62 (dt, *J* = 1.8, 0.8 Hz, 1H), 7.35 (dt, *J* = 8.6, 0.9 Hz, 1H), 7.28 (dd, *J* = 8.6, 1.8 Hz, 1H), 7.18 (dd, *J* = 3.2, 2.4 Hz, 1H), 6.53 (ddd, *J* = 3.0, 2.0, 0.8 Hz, 1H), 2.16 – 2.08 (m, 3H), 2.00 (d, *J* = 2.8 Hz, 6H), 1.79 (m, 6H). The procedure of **P2** synthesis: A dry, argon flushed flask equipped with a magnetic stirring bar was charged with **P1** (502 mg, 2.0 mmol, 1.0 equiv) in THF (10 mL), freshly distilled oxalyl chloride (279 mg, 2.2 mmol, 1.1 equiv) in anhydrous THF (2 mL) was added dropwise over 10 min at 0°C. An orange precipitate formed. Sat. aq NaHCO_3_ (5 mL) was then added with caution, and the mixture was heated at reflux for 30 min, then cooled and acidified with 10% HCl; this resulted in the precipitation of the substituted indole-3-glyoxalic acid, which was filtered and dried. Hydrazine hydrate (320 mg, 10 mmol, 5 equiv) was added to a solution of substituted indole-3-glyoxalic acid in 2-methoxyethanol (5 mL). The temperature of the mixture was increased to 60°C and NaOMe (1.08 g, 20 mmol, 10 equiv) was added portion-wise. The mixture was slowly heated at 150°C, whereby MeOH, H_2_O, hydrazine, and part of the solvent were evaporated. The mixture was kept at 150°C for 1 h, cooled, and poured onto crushed ice. The aqueous layer was extracted with EtOAc and acidified with concentrated HCl at 0°C. The oil that formed was extracted with EtOAc. The EtOAc solution was washed with H_2_O, dried over MgSO_4_, and evaporated in vacuo. The crude product obtained was further purified by the preparation HPLC to afford the desired pure product **P2** (white solid, 216 mg, 35% yield). ^1^H NMR (400 MHz, Chloroform-*d*) δ 8.03 (br s, 1H), 7.55 (d, *J* = 1.6 Hz, 1H), 7.30 (d, *J* = 1.4 Hz, 2H), 7.12 (d, *J* = 2.4 Hz, 1H), 3.82 (s, 2H), 2.16 – 2.08 (m, 3H), 2.00 (d, *J* = 2.8 Hz, 6H), 1.80 (m, 6H).

**Figure S2.**
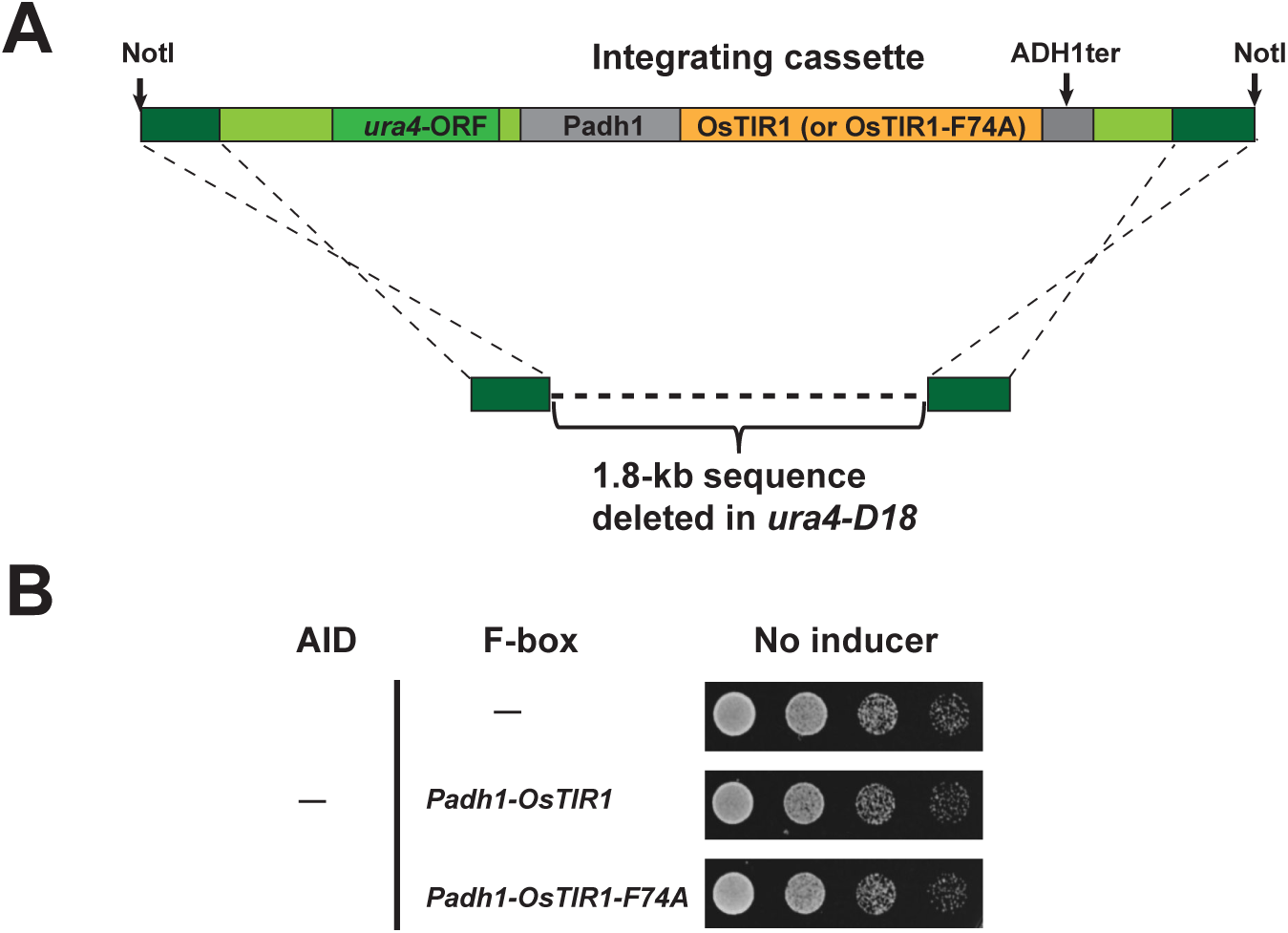
Integrating cassettes expressing OsTIR1 and OsTIR1-F74A from the *Padh1* promoter and the lack of toxicity of OsTIR1 and OsTIR1-F74A expressed from the *Padh1* promoter. **A.** Schematic showing the integrating cassette *Padh1-OsTIR1* (or *Padh1-OsTIR1-F74A*) released from plasmid pDB5050 (or pDB4695) by NotI digestion and the integration of the cassette through ends-out recombination at the *ura4* locus of a *ura4-D18* strain. Sequences from the *ura4* locus of *S. pombe* genome are depicted in green color of different shades. The homology arm sequences are depicted in dark green. The *ura4* coding sequence is depicted in medium green. Sequences between a homology arm and the *ura4* coding sequence are depicted in light green. *S. pombe Padh1* promoter and the terminator of *S. cerevisiae ADH1* gene (*ADH1ter*) are depicted in grey. The codon-optimized coding sequence of *OsTIR1* (or *OsTIR1-F74A*) is depicted in yellow. **B.** Spot assay showed that the *Padh1-OsTIR1* cassette and the *Padh1-OsTIR1-F74A* cassette, when integrated into the genome, did not cause any obvious growth phenotype in the absence of AID degron and inducer.

**Figure S3.**
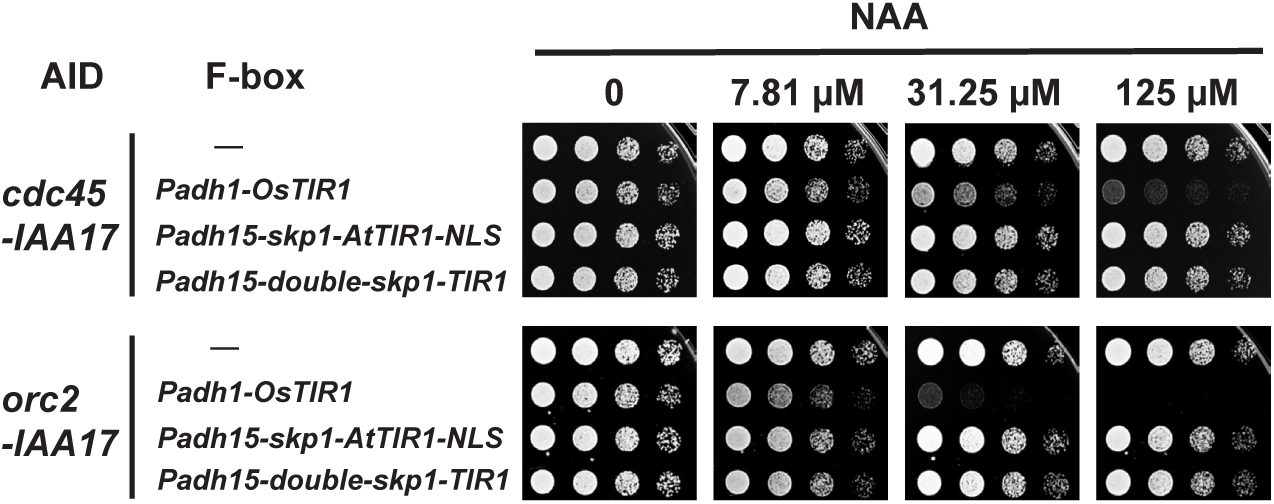
*Padh1*-expressed OsTIR1 is more effective than the F-box proteins used in the Kanke system for disrupting the functions of *cdc45* and *orc2*. Spot assay showed that when combined with *cdc45-IAA17* or *orc2-IAA17*, the *Padh1-OsTIR1* cassette was more effective at causing NAA-dependent growth inhibition than the F-box-protein-expressing cassettes of the Kanke system.

**Figure S4.**
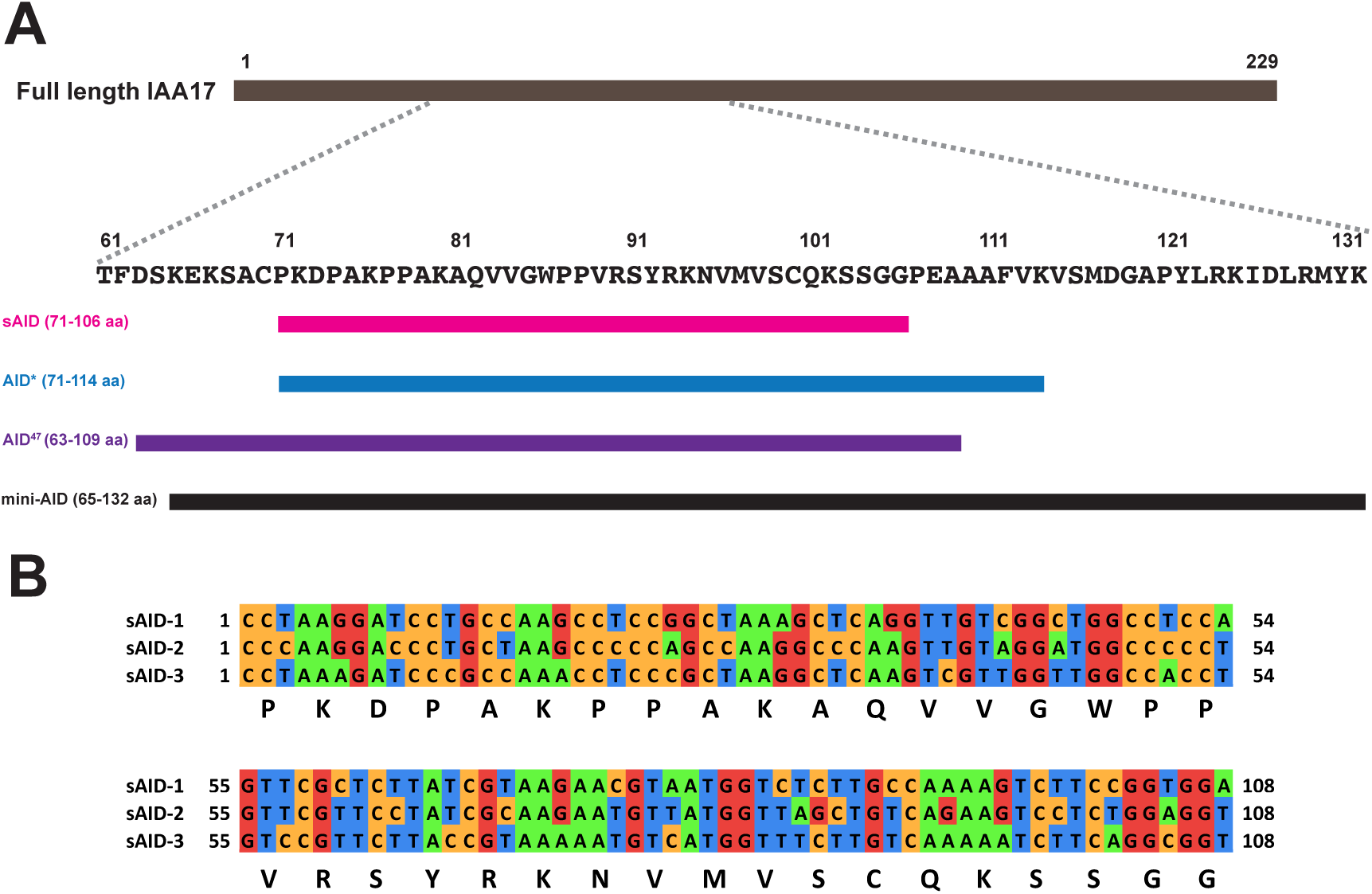
The design of the 3×sAID degron. A. Sequence of amino acids 61-132 of IAA17 and bars depicting four AID variants (sAID, AID*, AID^47^, and mini-AID) that correspond to parts of this region of IAA17. B. Sequence alignment of the nucleotide sequences encoding the three copies of sAID in the 3×sAID degron. Synonymous variations were intentionally introduced to reduce the chance of recombination between copies.

**Figure S5.**
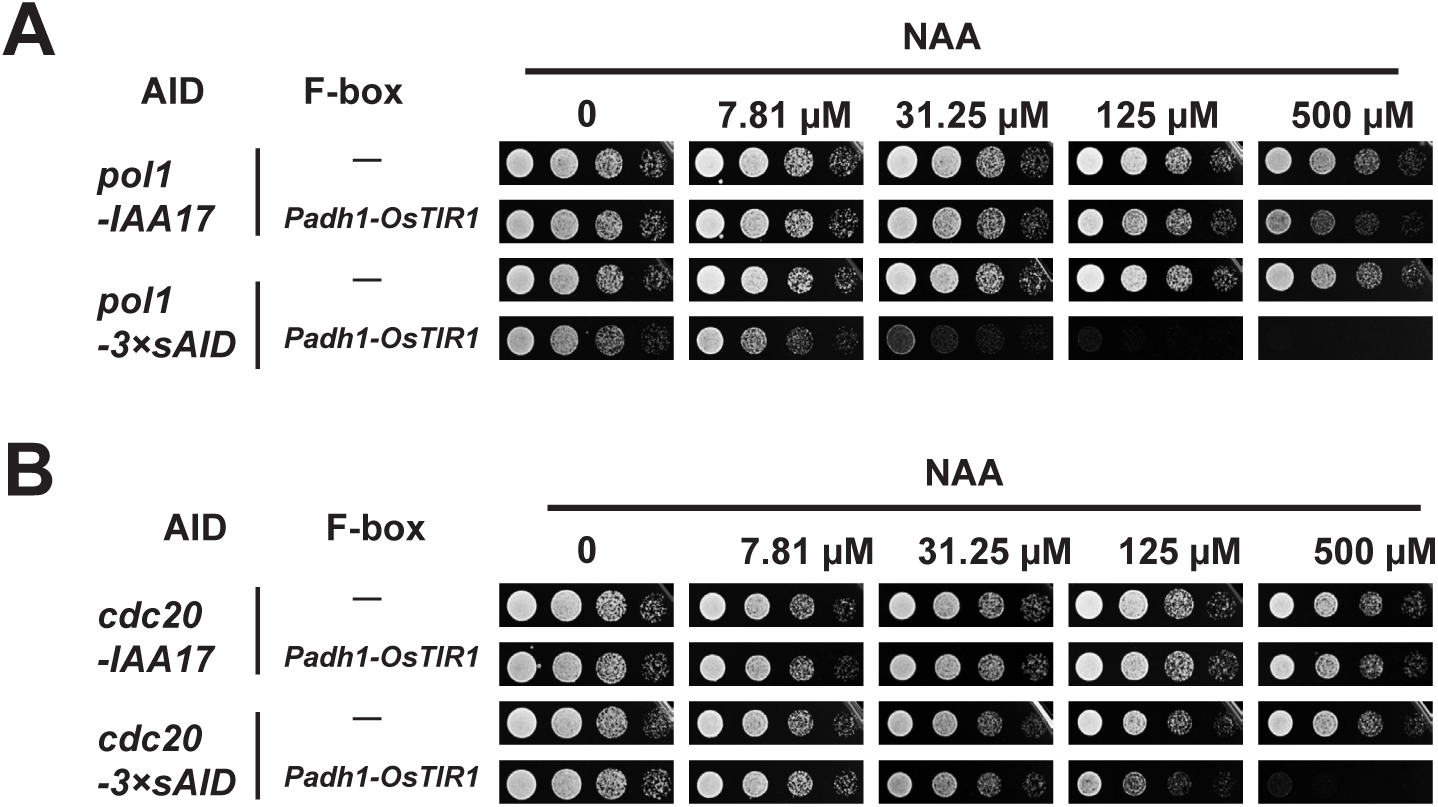
The 3×sAID degron is more effective than the full-length IAA17 for disrupting the functions of *pol1* and *cdc20*. A. Spot assay showed that when targeting the essential gene *pol1*, the 3×sAID degron was more effective at causing NAA-dependent growth inhibition than the full-length IAA17. B. Spot assay showed that when targeting the essential gene *cdc20*, the 3×sAID degron was more effective at causing NAA-dependent growth inhibition than the full-length IAA17.

**Figure S6.**
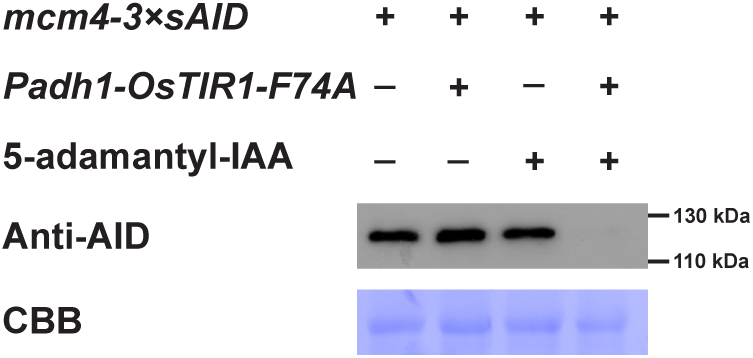
Complete protein depletion can be achieved within 15 min upon the addition of 5-adamantyl IAA. Two *mcm4-3×sAID* strains, one carrying *Padh1-OsTIR1-F74A* and one not, were cultured at 30°C in PMG media. The culture of each strain was split into two subcultures; one subculture was left untreated while 100 nM of 5-adamantyl-IAA was added to the other culture. After incubation for 15 minutes, cells were collected and lysed. Proteins in whole-cell extracts were separated on a 10% SDS-PAGE gel and analyzed by immunoblotting with an anti-mini-AID monoclonal antibody (MBL M214-3). Coomassie brilliant blue (CBB) staining of PVDF membrane was used as loading control.

**Table S1.**
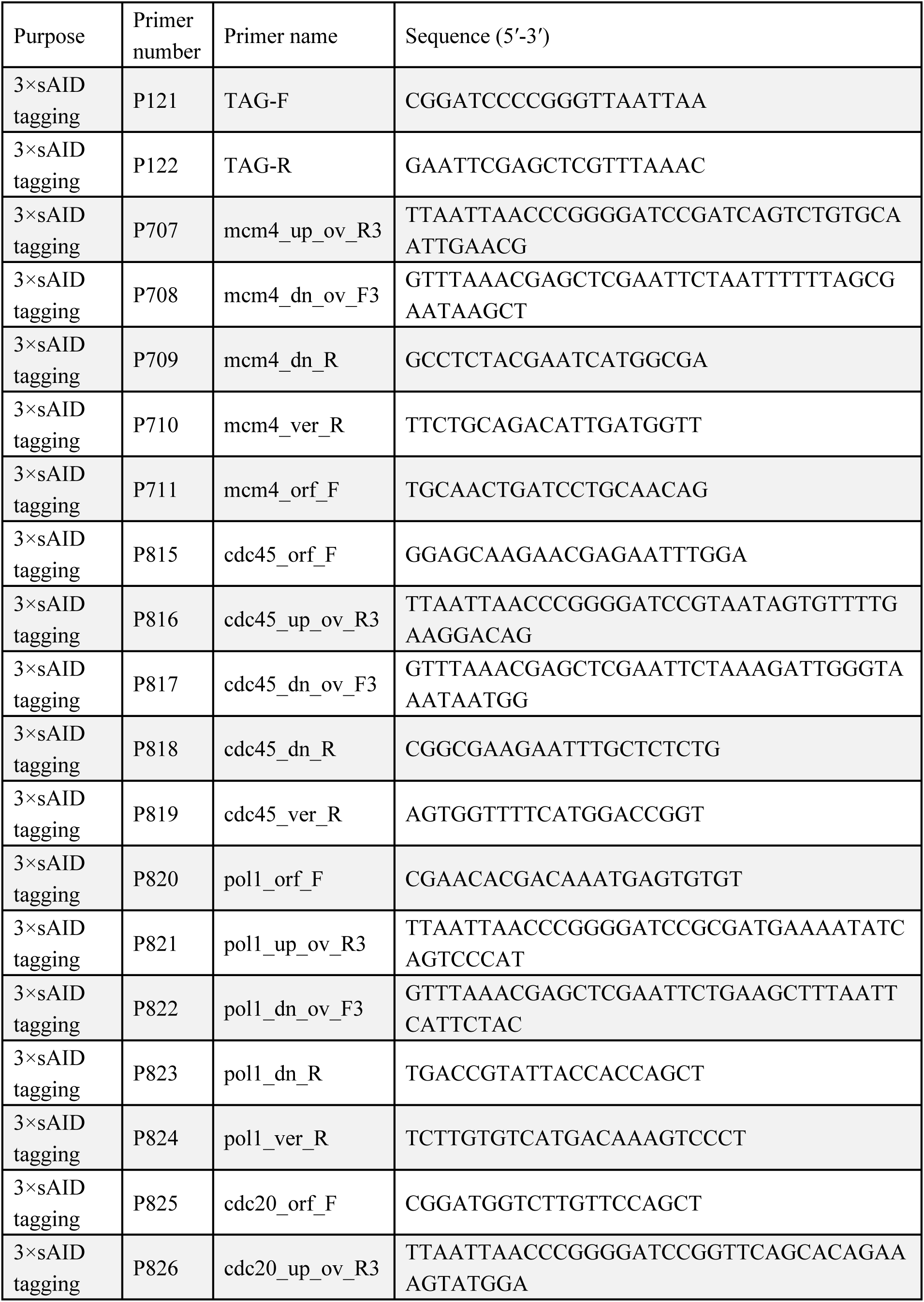
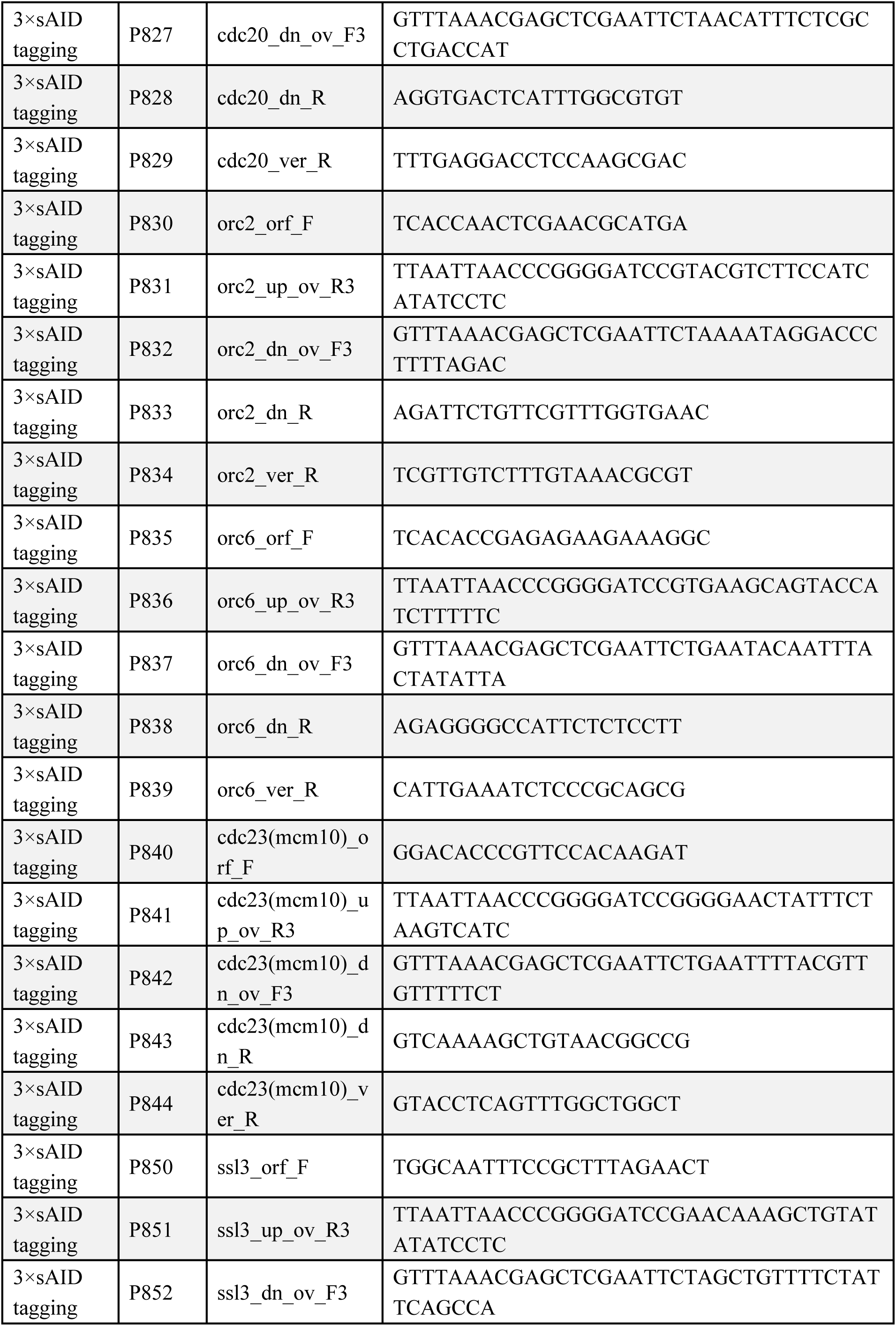
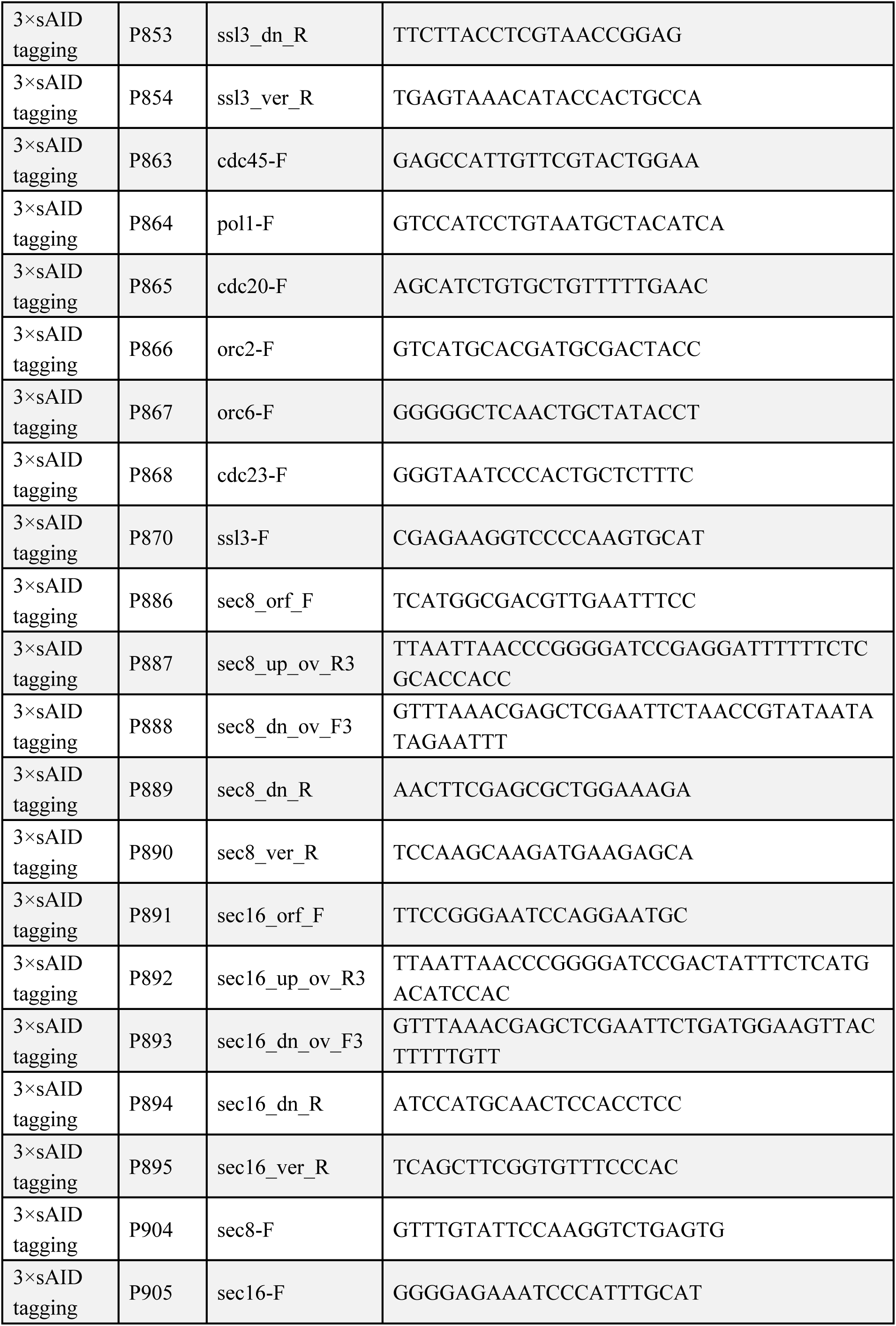
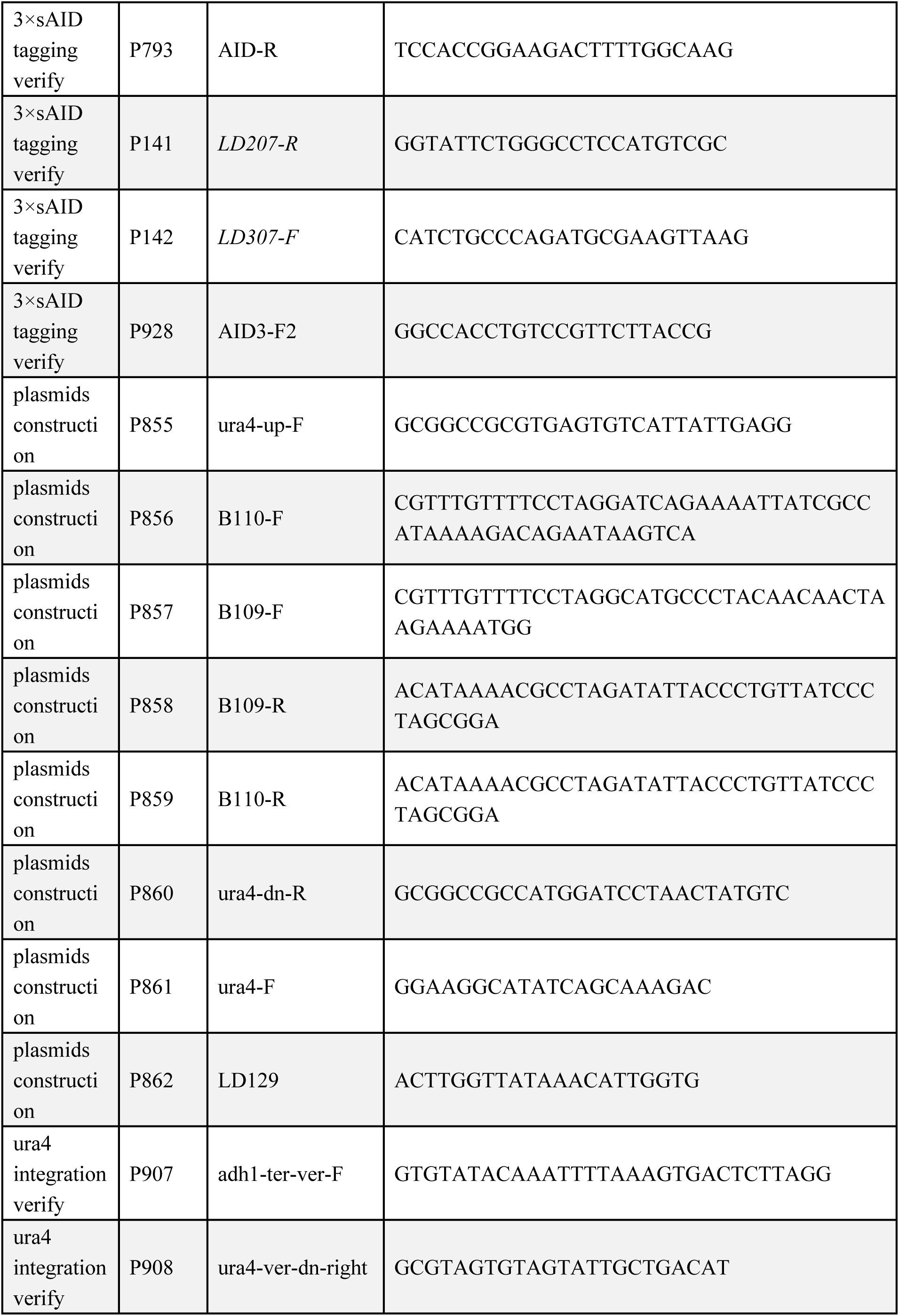
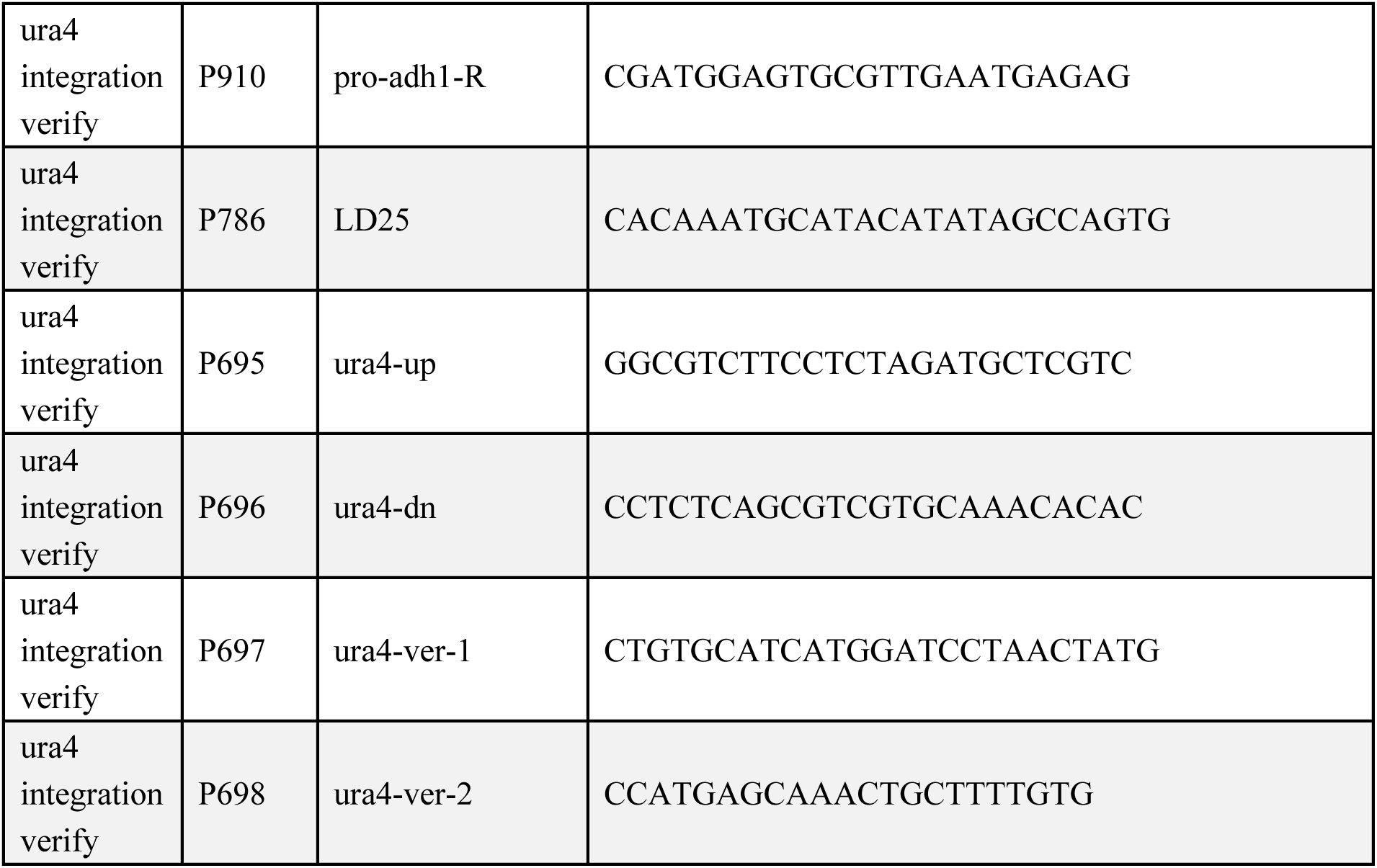
Primers used in this study.

**Table S2.**
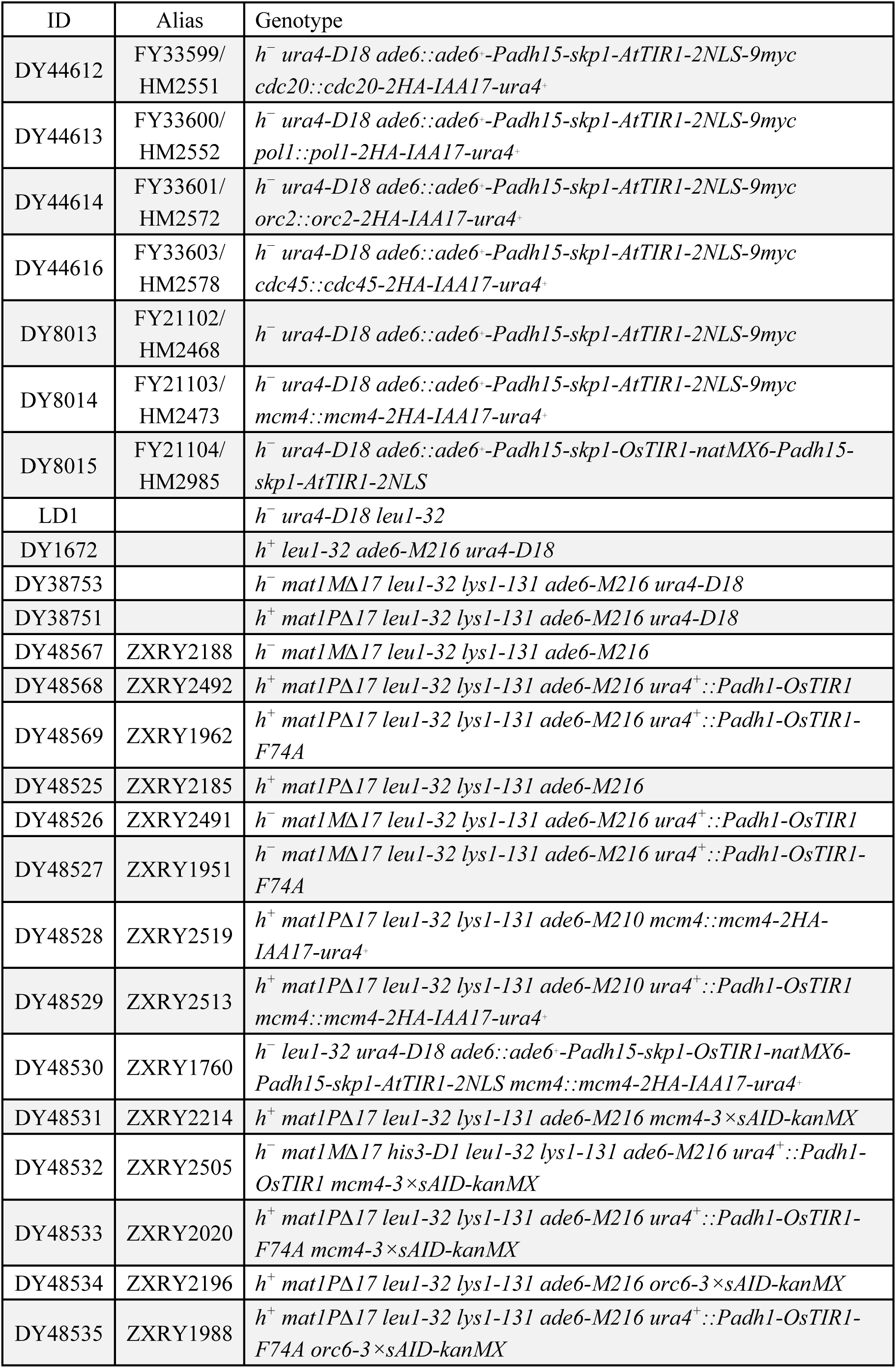
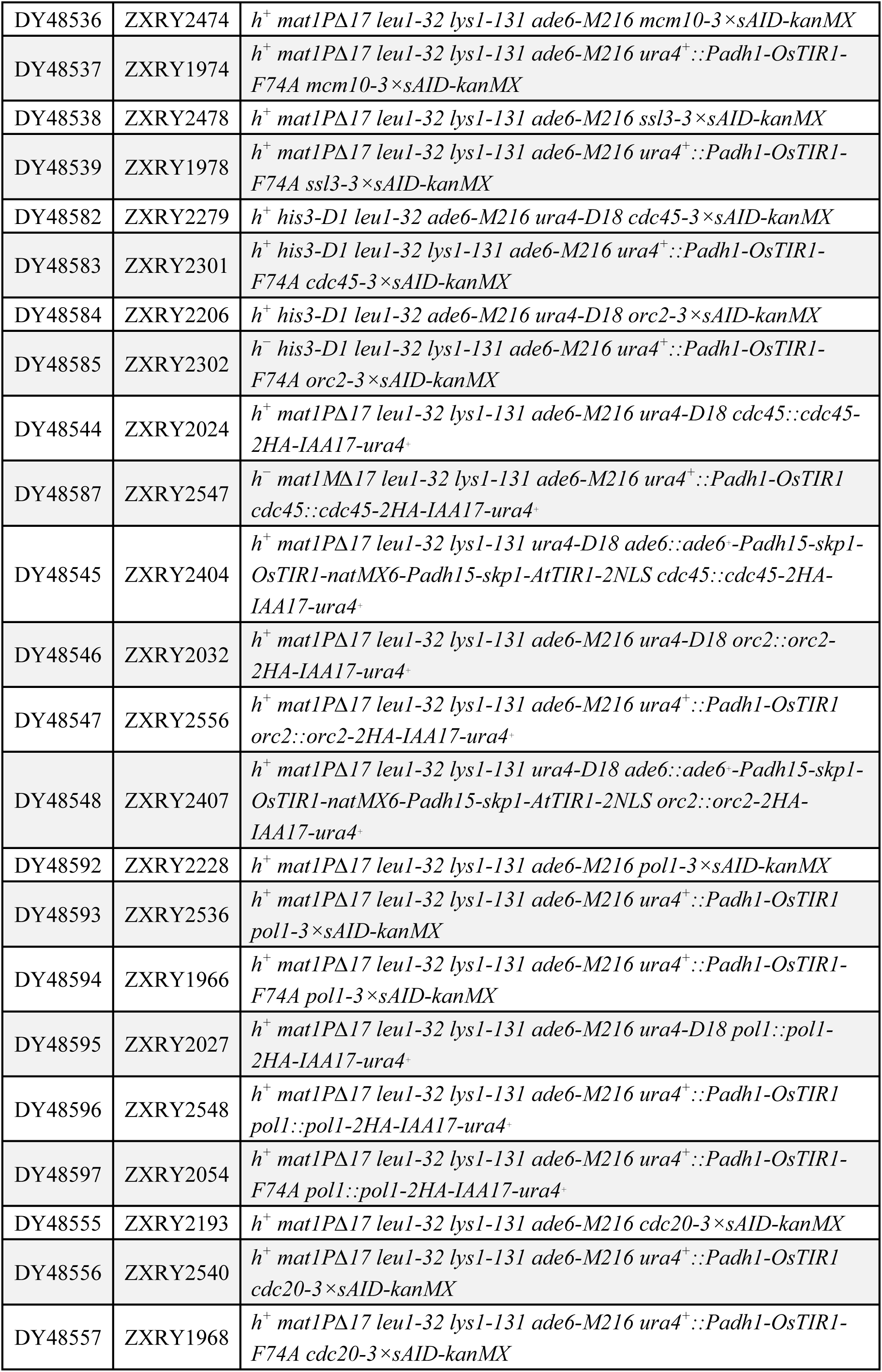
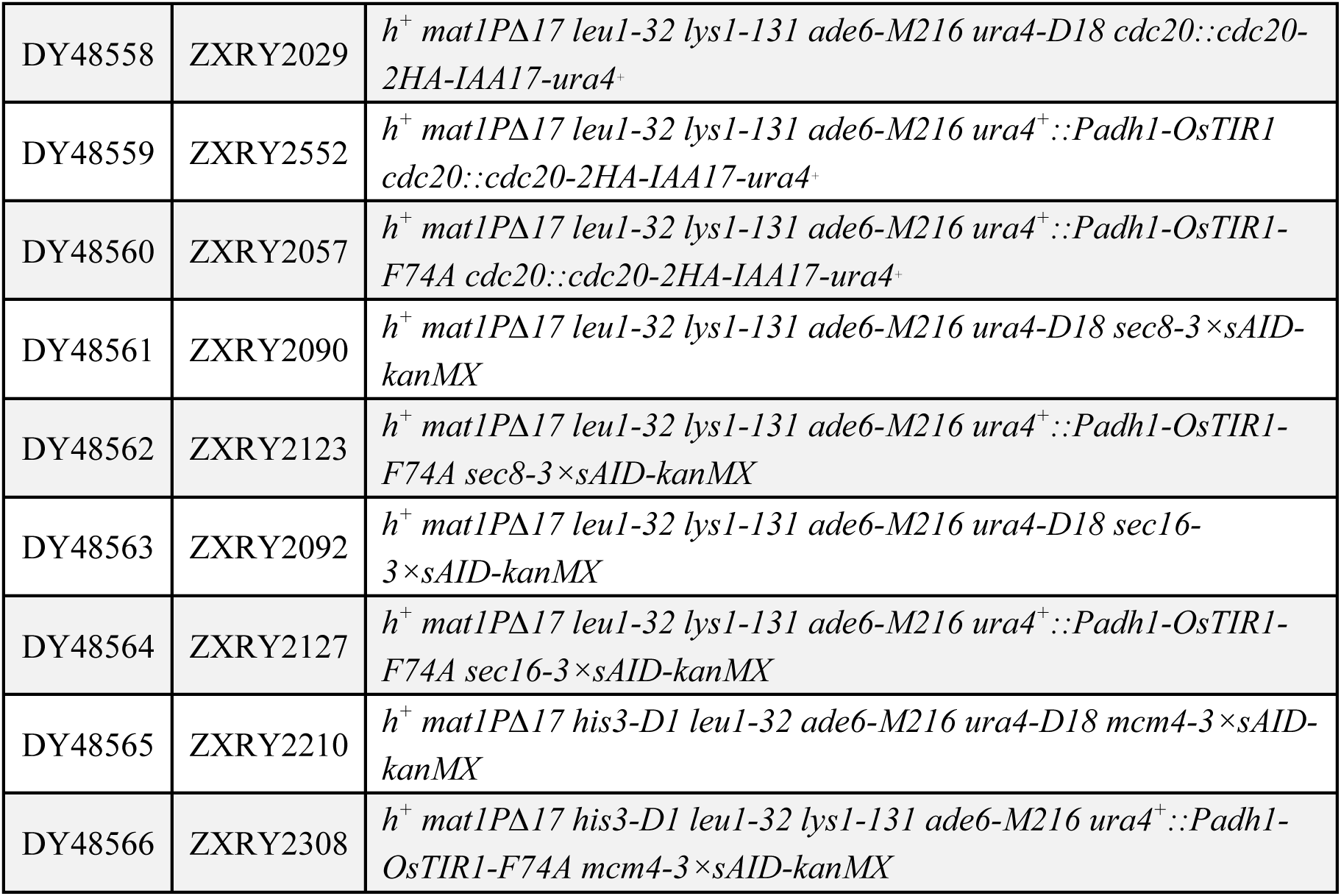
Fission yeast strains used in this study.

The following table is provided separately as an Excel file.

Table S3. RNA-seq results.

## References

Bahler, J., J.Q. Wu, M.S. Longtine, N.G. Shah, A. McKenzie, III et al., 1998 Heterologous modules for efficient and versatile PCR-based gene targeting in Schizosaccharomyces pombe. Yeast 14 (10):943–951.

Basi, G., E. Schmid, and K. Maundrell, 1993 TATA box mutations in the Schizosaccharomyces pombe nmt1 promoter affect transcription efficiency but not the transcription start point or thiamine repressibility. Gene 123 (1):131–136.

Ben-Aroya, S., C. Coombes, T. Kwok, K.A. O’Donnell, J.D. Boeke et al., 2008 Toward a comprehensive temperature-sensitive mutant repository of the essential genes of Saccharomyces cerevisiae. Mol Cell 30 (2):248–258.

Boe, C.A., I. Garcia, C.C. Pai, J.R. Sharom, H.C. Skjolberg et al., 2008 Rapid regulation of protein activity in fission yeast. BMC Cell Biol 9:23.

Brosh, R., I. Hrynyk, J. Shen, A. Waghray, N. Zheng et al., 2016 A dual molecular analogue tuner for dissecting protein function in mammalian cells. Nat Commun 7:11742.

Chanclud, E., and J.B. Morel, 2016 Plant hormones: a fungal point of view. Mol Plant Pathol 17 (8):1289–1297.

Costa, E.A., K. Subramanian, J. Nunnari, and J.S. Weissman, 2018 Defining the physiological role of SRP in protein-targeting efficiency and specificity. Science 359 (6376):689–692.

Ding, L., D. Laor, R. Weisman, and S.L. Forsburg, 2014 Rapid regulation of nuclear proteins by rapamycin-induced translocation in fission yeast. Yeast 31 (7):253–264.

Dohmen, R.J., P. Wu, and A. Varshavsky, 1994 Heat-inducible degron: a method for constructing temperature-sensitive mutants. Science 263 (5151):1273–1276.

Forsburg, S.L., and N. Rhind, 2006 Basic methods for fission yeast. Yeast 23 (3):173–183.

Fu, S.F., J.Y. Wei, H.W. Chen, Y.Y. Liu, H.Y. Lu et al., 2015 Indole-3-acetic acid: A widespread physiological code in interactions of fungi with other organisms. Plant Signal Behav 10 (8):e1048052.

Guilinger, J.P., D.B. Thompson, and D.R. Liu, 2014 Fusion of catalytically inactive Cas9 to FokI nuclease improves the specificity of genome modification. Nat Biotechnol 32 (6):577–582.

Haruki, H., J. Nishikawa, and U.K. Laemmli, 2008 The anchor-away technique: rapid, conditional establishment of yeast mutant phenotypes. Mol Cell 31 (6):925–932.

Kamisaka, S., N. Yanagishima, and Y. Masuda, 1967 Effect of Auxin and Gibberellin on Sporulation in Yeast. Physiol Plant 20 (1):90–97.

Kanemaki, M., A. Sanchez-Diaz, A. Gambus, and K. Labib, 2003 Functional proteomic identification of DNA replication proteins by induced proteolysis in vivo. Nature 423 (6941):720–724.

Kanke, M., K. Nishimura, M. Kanemaki, T. Kakimoto, T.S. Takahashi et al., 2011 Auxin-inducible protein depletion system in fission yeast. BMC Cell Biol 12:8.

Kearsey, S.E., and J. Gregan, 2009 Using the DHFR heat-inducible degron for protein inactivation in Schizosaccharomyces pombe. Methods Mol Biol 521:483–492.

Kim, D., J.M. Paggi, C. Park, C. Bennett, and S.L. Salzberg, 2019 Graph-based genome alignment and genotyping with HISAT2 and HISAT-genotype. Nat Biotechnol 37 (8):907–915.

Kubota, T., K. Nishimura, M.T. Kanemaki, and A.D. Donaldson, 2013 The Elg1 replication factor C-like complex functions in PCNA unloading during DNA replication. Mol Cell 50 (2):273–280.

Li, S., X. Prasanna, V.T. Salo, I. Vattulainen, and E. Ikonen, 2019 An efficient auxin-inducible degron system with low basal degradation in human cells. Nat Methods 16 (9):866–869.

Liao, Y., G.K. Smyth, and W. Shi, 2014 featureCounts: an efficient general purpose program for assigning sequence reads to genomic features. Bioinformatics 30 (7):923–930.

Liu, Y.Y., H.W. Chen, and J.Y. Chou, 2016 Variation in Indole-3-Acetic Acid Production by Wild Saccharomyces cerevisiae and S. paradoxus Strains from Diverse Ecological Sources and Its Effect on Growth. PLoS One 11 (8):e0160524.

Longtine, M.S., A. McKenzie, 3rd, D.J. Demarini, N.G. Shah, A. Wach et al., 1998 Additional modules for versatile and economical PCR-based gene deletion and modification in Saccharomyces cerevisiae. Yeast 14 (10):953–961.

Love, M.I., W. Huber, and S. Anders, 2014 Moderated estimation of fold change and dispersion for RNA-seq data with DESeq2. Genome Biol 15 (12):550.

Lyne, R., G. Burns, J. Mata, C.J. Penkett, G. Rustici et al., 2003 Whole-genome microarrays of fission yeast: characteristics, accuracy, reproducibility, and processing of array data. BMC Genomics 4 (1):27.

Mendoza-Ochoa, G.I., J.D. Barrass, B.R. Terlouw, I.E. Maudlin, S. de Lucas et al., 2019 A fast and tuneable auxin-inducible degron for depletion of target proteins in budding yeast. Yeast 36 (1):75–81.

Mnaimneh, S., A.P. Davierwala, J. Haynes, J. Moffat, W.T. Peng et al., 2004 Exploration of essential gene functions via titratable promoter alleles. Cell 118 (1):31–44.

Morawska, M., and H.D. Ulrich, 2013 An expanded tool kit for the auxin-inducible degron system in budding yeast. Yeast 30 (9):341–351.

Mutte, S.K., H. Kato, C. Rothfels, M. Melkonian, G.K. Wong et al., 2018 Origin and evolution of the nuclear auxin response system. Elife 7.

Natsume, T., and M.T. Kanemaki, 2017 Conditional Degrons for Controlling Protein Expression at the Protein Level. Annu Rev Genet 51:83–102.

Natsume, T., T. Kiyomitsu, Y. Saga, and M.T. Kanemaki, 2016 Rapid Protein Depletion in Human Cells by Auxin-Inducible Degron Tagging with Short Homology Donors. Cell Rep 15 (1):210–218.

Nicastro, R., S. Raucci, A.H. Michel, M. Stumpe, G.M. Garcia Osuna et al., 2021 Indole-3-acetic acid is a physiological inhibitor of TORC1 in yeast. PLoS Genet 17 (3):e1009414.

Nishimura, K., T. Fukagawa, H. Takisawa, T. Kakimoto, and M. Kanemaki, 2009 An auxin-based degron system for the rapid depletion of proteins in nonplant cells. Nat Methods 6 (12):917–922.

Nishimura, K., and M.T. Kanemaki, 2014 Rapid Depletion of Budding Yeast Proteins via the Fusion of an Auxin-Inducible Degron (AID). Curr Protoc Cell Biol 64:20 29 21-16.

Nishimura, K., R. Yamada, S. Hagihara, R. Iwasaki, N. Uchida et al., 2020 A super-sensitive auxin-inducible degron system with an engineered auxin-TIR1 pair. Nucleic Acids Res 48 (18):e108.

Pai, C.C., J. Schnick, S.A. MacNeill, and S.E. Kearsey, 2012 Conditional inactivation of replication proteins in fission yeast using hormone-binding domains. Methods 57 (2):227–233.

Petit, S., Y. Duroc, V. Larue, C. Giglione, C. Leon et al., 2009 Structure-activity relationship analysis of the peptide deformylase inhibitor 5-bromo-1H-indole-3-acetohydroxamic acid. ChemMedChem 4 (2):261–275.

Prusty, R., P. Grisafi, and G.R. Fink, 2004 The plant hormone indoleacetic acid induces invasive growth in Saccharomyces cerevisiae. Proc Natl Acad Sci U S A 101 (12):4153–4157.

Rajagopalan, S., Z. Liling, J. Liu, and M. Balasubramanian, 2004 The N-degron approach to create temperature-sensitive mutants in Schizosaccharomyces pombe. Methods 33 (3):206–212.

Rao, R.P., A. Hunter, O. Kashpur, and J. Normanly, 2010 Aberrant synthesis of indole-3-acetic acid in Saccharomyces cerevisiae triggers morphogenic transition, a virulence trait of pathogenic fungi. Genetics 185 (1):211–220.

Samann, C., V. Dhayalan, P.R. Schreiner, and P. Knochel, 2014 Synthesis of substituted adamantylzinc reagents using a Mg-insertion in the presence of ZnCl(2) and further functionalizations. Org Lett 16 (9):2418–2421.

Sathyan, K.M., B.D. McKenna, W.D. Anderson, F.M. Duarte, L. Core et al., 2019 An improved auxin-inducible degron system preserves native protein levels and enables rapid and specific protein depletion. Genes Dev 33 (19-20):1441–1455.

Snyder, N.A., A. Kim, L. Kester, A.N. Gale, C. Studer et al., 2019 Auxin-Inducible Depletion of the Essentialome Suggests Inhibition of TORC1 by Auxins and Inhibition of Vrg4 by SDZ 90-215, a Natural Antifungal Cyclopeptide. G3 (Bethesda) 9 (3):829–840.

Tan, X., L.I. Calderon-Villalobos, M. Sharon, C. Zheng, C.V. Robinson et al., 2007 Mechanism of auxin perception by the TIR1 ubiquitin ligase. Nature 446 (7136):640–645.

Tang, X., J. Huang, A. Padmanabhan, K. Bakka, Y. Bao et al., 2011 Marker reconstitution mutagenesis: a simple and efficient reverse genetic approach. Yeast 28 (3):205–212.

Texier, P., M. Coddeville, P. Bordes, and P. Genevaux, 2018 A useful gene cassette for conditional knock-down of essential genes by targeted promoter replacement in Mycobacteria. Biotechniques 65 (3):159–162.

Timney, B.L., B. Raveh, R. Mironska, J.M. Trivedi, S.J. Kim et al., 2016 Simple rules for passive diffusion through the nuclear pore complex. J Cell Biol 215 (1):57–76.

Torii, K.U., S. Hagihara, N. Uchida, and K. Takahashi, 2018 Harnessing synthetic chemistry to probe and hijack auxin signaling. New Phytol 220 (2):417–424.

Uchida, N., K. Takahashi, R. Iwasaki, R. Yamada, M. Yoshimura et al., 2018 Chemical hijacking of auxin signaling with an engineered auxin-TIR1 pair. Nat Chem Biol 14 (3):299–305.

Ulrich, H.D., and A.A. Davies, 2009 In vivo detection and characterization of sumoylation targets in Saccharomyces cerevisiae. Methods Mol Biol 497:81–103.

Watson, A.T., Y. Daigaku, S. Mohebi, T.J. Etheridge, C. Chahwan et al., 2013 Optimisation of the Schizosaccharomyces pombe urg1 expression system. PLoS One 8 (12):e83800.

Watson, A.T., S. Hassell-Hart, J. Spencer, and A.M. Carr, 2021 Rice (Oryza sativa) TIR1 and 5’adamantyl-IAA Significantly Improve the Auxin-Inducible Degron System in Schizosaccharomyces pombe. Genes (Basel) 12 (6):886.

Yamada, R., K. Murai, N. Uchida, K. Takahashi, R. Iwasaki et al., 2018 A Super Strong Engineered Auxin-TIR1 Pair. Plant Cell Physiol 59 (8):1538–1544.

Yanagishima, N., and Y. Masuda, 1965 Further Studies on RNA in Relation to Auxin-Induced Cell Elongation in Yeast. Physiol Plant 18 (3):586–591.

Yesbolatova, A., T. Natsume, K.I. Hayashi, and M.T. Kanemaki, 2019 Generation of conditional auxin-inducible degron (AID) cells and tight control of degron-fused proteins using the degradation inhibitor auxinole. Methods 164–165:73-80.

Yesbolatova, A., Y. Saito, N. Kitamoto, H. Makino-Itou, R. Ajima et al., 2020 The auxin-inducible degron 2 technology provides sharp degradation control in yeast, mammalian cells, and mice. Nat Commun 11 (1):5701.

Zhang, L., J.D. Ward, Z. Cheng, and A.F. Dernburg, 2015 The auxin-inducible degradation (AID) system enables versatile conditional protein depletion in C. elegans. Development 142 (24):4374–4384.

